# The circulating cell-free DNA landscape in sepsis is dominated by impaired liver clearance

**DOI:** 10.1101/2025.02.17.638622

**Authors:** Kiki Cano-Gamez, Patrick Maclean, Masato Inoue, Sakineh Hussainy, Elisabeth Foss, Chloe Wainwright, Hanyu Qin, Stuart McKechnie, Chun-Xiao Song, Julian C. Knight

## Abstract

Circulating cell-free DNA (cfDNA) is a promising molecular biomarker. However, its utility in severe infection remains poorly understood. Here, we isolated cfDNA from sepsis patients and controls, demonstrating a 41-fold increase in the amount of cfDNA in circulation during disease. We used sequencing to reconstruct cfDNA methylomes, fragmentation profiles, and nucleosome footprints across 56 samples. We observed no difference in cfDNA composition between patients and controls, challenging the idea that cfDNA increases due to higher immune cell death during sepsis. Instead, we suggest that liver dysfunction prevents efficient clearance of cfDNA during disease. This was supported by fragmentation and end-motif patterns, both of which showed evidence of cfDNA being exposed to circulating nucleases for a prolonged period, proportionally to the extent of liver dysfunction. Variation in cfDNA of megakaryocyte-erythroid progenitor origin was also a significant contributor in sepsis, increasing over time. Moreover, we showed that cfDNA retains nucleosome footprints with cell type-specific gene activity information. We developed a novel approach to study nucleosome phasing that successfully recovers tissue-specific signatures. By combining this with single-cell data, we demonstrated that sepsis patients with liver dysfunction have higher amounts of cfDNA derived from Kupffer cells and the liver parenchyma. In conclusion, we present the first high-throughput multi-modal study of cfDNA during sepsis, which will serve as a reference point for future studies on the role of this biomarker in critical illness.

## Introduction

Circulating cell-free DNA (cfDNA) comprises a collection of DNA fragments which exist outside their cellular context and circulate freely in the bloodstream. These fragments are thought to originate from homeostatic cell death (e.g. apoptosis and efferocytosis) as dying cells release apoptotic bodies and chromatin fragments into the circulation for clearance by the mononuclear phagocyte system (1). The half-life of cfDNA is estimated to be between several minutes and 2 hours (1), which makes it ideal as a potential biomarker for real-time monitoring of cell death at remote tissues and organs. Circulating cfDNA has been widely used as a biomarker in pregnancy, organ transplantation, and cancer, where the mixture of genotypes in the cfDNA pool makes it possible to detect pregnancy complications (2,3), organ rejection (4–6), and mutated tumour-derived DNA (7–10) before clinical signs of illness are apparent. However, the utility of cfDNA in acute conditions like severe infection and critical illness remains underexplored.

The rapid and non-invasive nature of cfDNA profiling makes it particularly promising in the context of sepsis, the dysregulated host response to infection associated with life-threatening organ dysfunction (11,12). Sepsis is one of the leading causes of death worldwide, with 11 million deaths estimated to be caused by this clinical syndrome every year (13). Biomarkers to enable earlier and more accurate detection of organ failure in sepsis would facilitate clinical decision-making and timely therapeutic intervention, helping address the burden of this devastating condition. In addition, sepsis is a highly heterogeneous syndrome with a variety of potential disease mechanisms operating in different patients (14–20). Consequently, monitoring of cfDNA could help elucidate the mechanisms underlying disease (e.g. by tracking immune cell function or identifying abnormal cell death events like endothelial damage and necrosis), thus improving diagnostic precision and enabling patient stratification (14).

Previous studies have demonstrated that cfDNA is significantly increased during sepsis and have suggested this relates to disease severity (21,22). For example, cfDNA concentration in serum is associated with sepsis mortality (23) and higher levels of circulating mitochondrial DNA (mtDNA) predict poor prognosis (24). However, the tissues of origin of this circulating DNA and how it may be cleared from the blood are not known, which complicates the interpretation of these findings.

Recent advances in experimental techniques to assay the epigenome (25,9,26–30) and computational tools to deconvolute bulk signals into cell type-specific contributions (6,31–34) now make it possible to infer which tissues contribute material to the circulation based on the presence of epigenetic marks in cfDNA (e.g. DNA methylation or chromatin modifications). A recent study profiled DNA methylation to better understand the composition of cfDNA in several conditions, including organ transplantation, cancer, and sepsis (6). Observations from this study suggest that sepsis may cause a sharp increase in the amount of cfDNA derived from neutrophils, which supports the known role of neutrophil extracellular traps (NETs) in this disease (35–37). Moreover, cfDNA composition was shown to reflect instances of organ failure (6). Despite these advances, an in-depth characterization of the circulating cfDNA landscape (the *circulome*) in sepsis is needed to fully understand its clinical potential.

Here, we present the first large-scale high-throughput study of the circulome during acute sepsis. By profiling cfDNA methylation, fragmentation patterns, end-motif sequences, and nucleosome positioning signatures across hospitalised sepsis patients, we present an in-depth characterisation of the tissues of origin of cfDNA and identify potential mechanisms driving cfDNA accumulation. Our observations confirm that cfDNA increases during acute sepsis but we observe no major differences in cfDNA composition between sepsis patients and controls, suggesting that cfDNA accumulation is not primarily driven by cell type-specific cell death. Instead, our observations are compatible with cfDNA accumulating due to impaired liver clearance, a hypothesis supported by both fragmentation patterns and end-motif frequencies. Finally, we show that analysis of methylation and nucleosome positioning in cfDNA could enable development of novel biomarkers for monitoring liver function, organ failure, and immune cell turnover during sepsis that could help monitor disease progression and detect adverse events during critical illness.

## Results

### Levels of cfDNA in circulation reflect sepsis severity

To assess the utility of cfDNA in sepsis, we isolated cfDNA from 86 plasma samples obtained from 46 patients admitted to hospital with sepsis and serially sampled as part of the Sepsis Immunomics study together with 12 healthy controls (**Methods**). In line with previous literature, we found that the cfDNA concentration was dramatically higher in sepsis patients compared to healthy controls, with an average 41.2-fold increase (**Figure 1A**). When stratifying patients by their required level of care, we found a significant increase in cfDNA in patients requiring higher care levels, with patients in the intensive care unit (ICU) showing concentrations approximately 10-fold higher than patients in the emergency department (ED) and general medical ward (**Figure 1A**).

**Figure 1.**
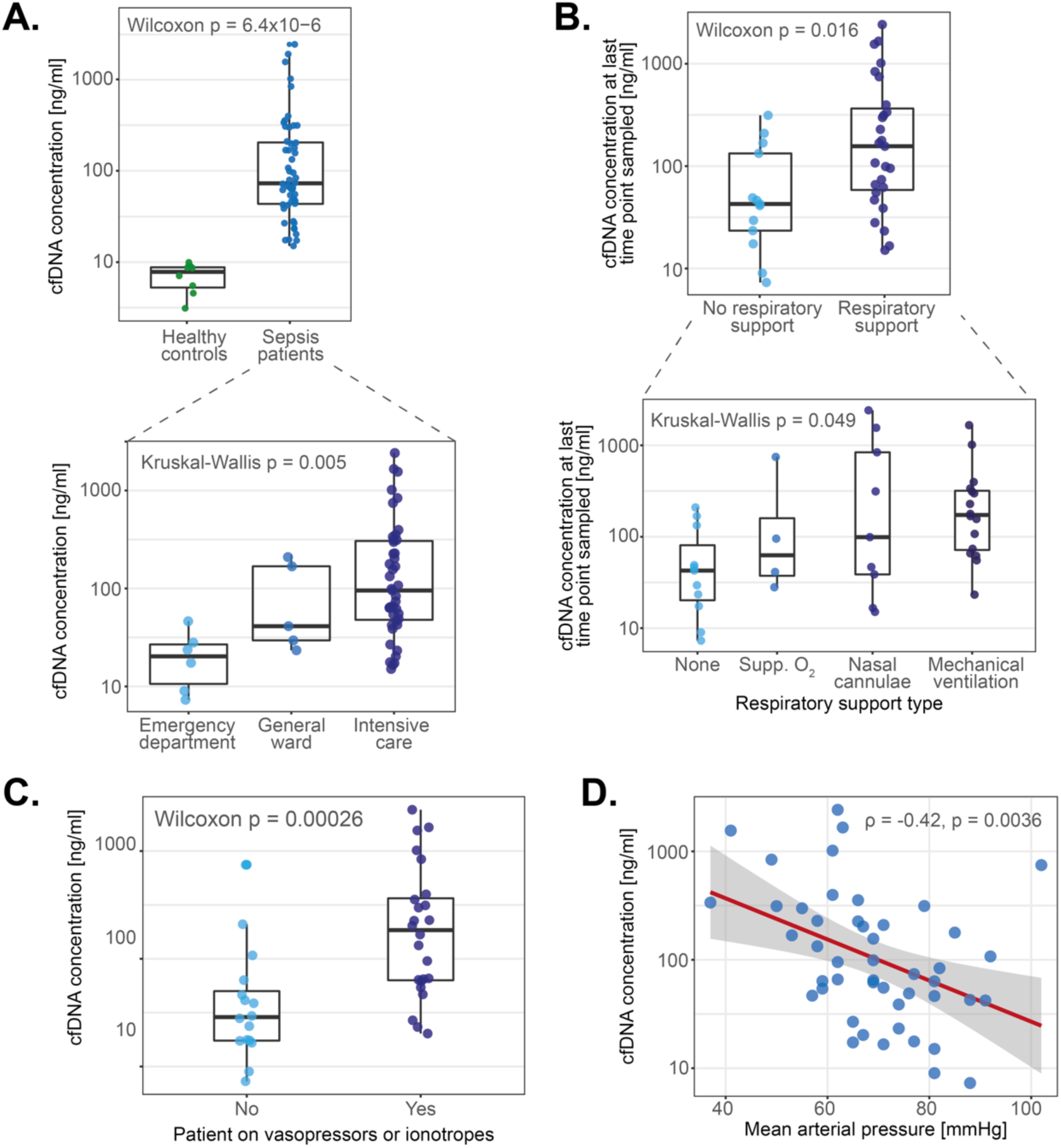
Levels of cfDNA in circulation reflect sepsis severity. **A.** cfDNA concentration in the plasma of sepsis patients and healthy controls (upper panel), and in sepsis patients within different hospital care settings (lower panel). P values were derived using a Wilcoxon Rank Sum test. Box plots represent median and IQRs cfDNA concentration. **B.** cfDNA concentration in sepsis patients stratified by level of respiratory support. Categories are ordered by increasing invasiveness. P values were derived using a Wilcoxon Rank Sum test. Box plots represent median and IQRs cfDNA concentration. **C.** cfDNA concentration in sepsis patients stratified by requirement for treatment with vasopressors or inotropes. P values were derived using a Wilcoxon Rank Sum test. Box plots represent median and IQRs cfDNA concentration. **D.** Correlation between mean arterial pressure (X axis) and cfDNA concentration (Y axis). P values and correlation estimates were computed with a Pearson correlation test.

To confirm the association between cfDNA concentration and illness severity, we further stratified patients by their required level of respiratory support, as well as by the presence of cardiovascular dysfunction requiring treatment (e.g. hypotension or shock). Cell-free DNA concentration increased gradually for patients requiring progressively more invasive forms of respiratory support (**Figure 1B**), suggesting it is related to the extent of respiratory failure. Moreover, cfDNA levels were significantly higher in patients requiring vasopressor or inotrope treatment (**Figure 1C**). This suggested a relationship between hypotension and cfDNA, as confirmed by a significant negative correlation between cfDNA concentration and mean arterial pressure (MAP) measured on the day of sampling (**Figure 1D**). Taken together, these observations demonstrate that cfDNA levels are reflective of illness severity, thus motivating further study on the biology and diagnostic utility of this molecular compartment.

### An approach for multi-modal profiling of circulating cfDNA

We next asked why cfDNA increases during sepsis. Based on our current understanding of sepsis pathophysiology (12), we identified a set of mechanisms which could explain the observed cfDNA accumulation. We grouped these into two: 1) mechanisms which increase cfDNA release, and 2) mechanisms which reduce cfDNA clearance (**Figure 2A**). The first group comprised cell death concomitant to the host immune response (e.g. NETosis, pyroptosis, necrosis), damage of the vascular endothelium, and organ failure. In contrast, the second group included hepatic clearance by Kupffer cells and renal filtration (25). We reasoned that the contribution of these competing mechanisms could be distinguished based on cfDNA composition, which motivated us to leverage the information contained in cfDNA methylation, fragmentation, and nucleosome positioning to identify the likely tissues of origin of cfDNA.

**Figure 2.**
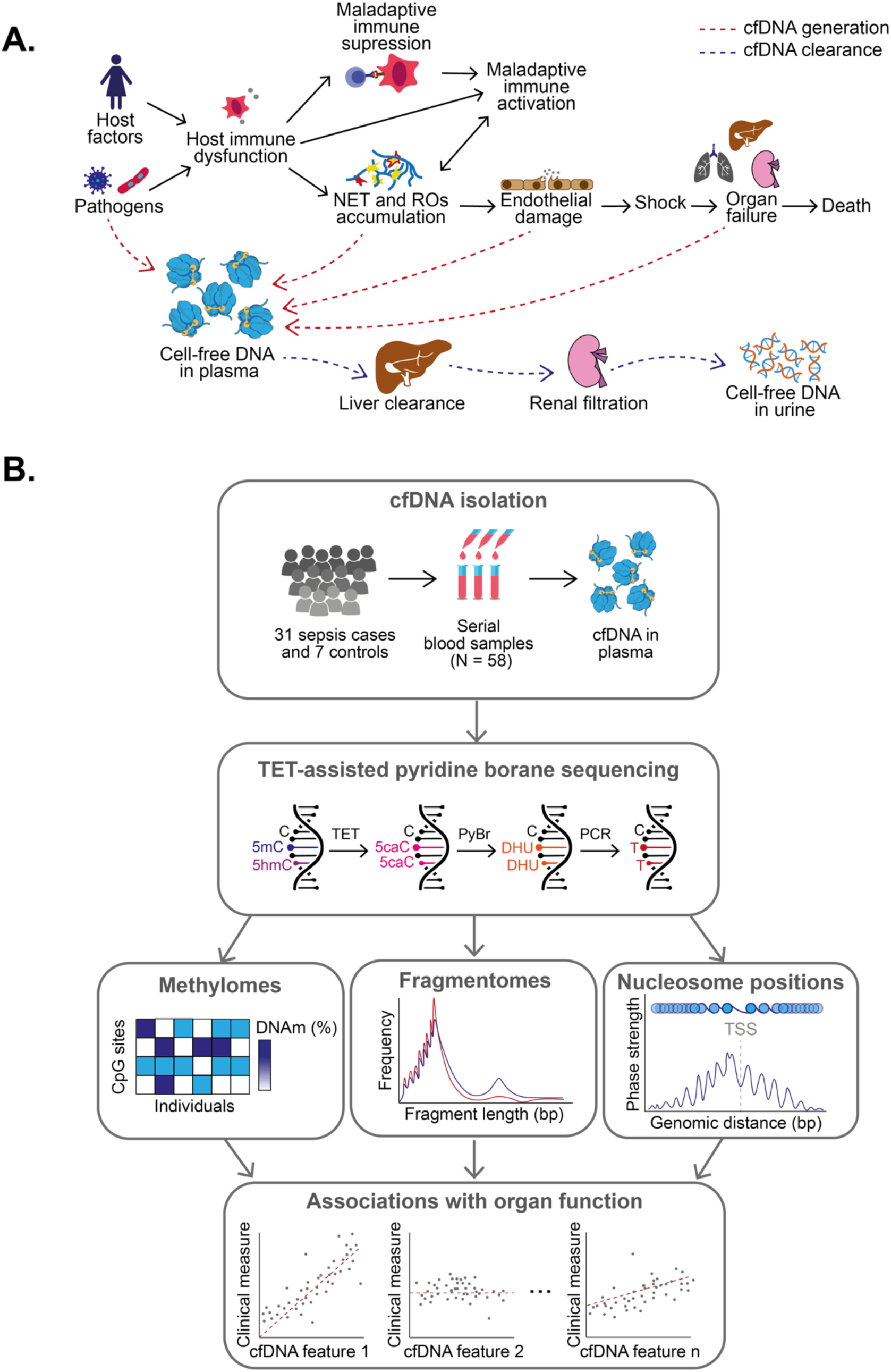
An experimental approach for multi-modal profiling of circulating cfDNA. **A.** Schematic of possible mechanisms contributing and clearing DNA from circulation. Solid arrows represent putative causal relationships between disease processes. Dashed red and blue arrows represent possible mechanisms of cfDNA production and clearance during sepsis, respectively. **B.** Experimental approach followed in this study. Blood samples were collected from sepsis patients and used for cfDNA isolation. Cell-free DNA was subsequently used for library preparation and sequencing with the TAPS protocol, which enabled simultaneous characterisation of its methylation status, fragmentation patterns, and evidence of nucleosome positioning. This information was then related back to clinical measures of illness severity and used to infer the likely tissues of origin of cfDNA.

We used TET-assisted pyridine borane sequencing (TAPS), a method which enables reliable genome-wide profiling of DNA methylation (9,25). In contrast to techniques like bisulfite-sequencing (BS-seq) (38,39) or enzymatic methyl-sequencing (EM-seq) (27), TAPS is ideally suited to low DNA input and its direct conversion of methylated cytosines does not significantly reduce sequence diversity, enabling efficient bioinformatic analysis of sequence-based DNA features. We used TAPS to profile DNA methylation in 58 cfDNA samples from 31 sepsis patients and 7 healthy controls, of which 56 passed our QC filters (**Methods**). Demographics and sample characteristics for this cohort are shown in **Figure S1**. We leveraged this data to study three layers of information: 1) the genome-wide cfDNA methylome, 2) the cfDNA fragmentation landscape (i.e. fragment size distributions and end-motif frequencies), and 3) nucleosome positioning at gene regions (**Figure 2B**). We used these complementary data modalities to infer the likely tissues of origin of cfDNA and to assess its potential utility as a biomarker for sepsis monitoring.

### Variation in the cfDNA methylome reflects organ function and disease processes

We first assessed the quality of our methylation data comprising all patients and controls. We confirmed that TAPS could successfully identify methylated CpG sites (i.e. 5mC + 5hmC) at 93% sensitivity with only a 0.35% false positive rate based on the rate of 5mC>A conversion of fully methylated and fully unmethylated spike-in controls (**Figure 3A**). This gave us confidence that our data was high quality. We proceeded to analyse a set of 19,288,064 autosomal CpG sites reliably detected across sample groups (**Methods**). To maximise power, we collapsed CpG sites into units based on genomic proximity by segmenting the genome into non-overlapping 1-kb tiles (**Methods**). In agreement with previous literature (40), this revealed two types of genomic regions: those with average methylation of approximately 0% (hypomethylated) and those with a mean methylation of 75% (hypermethylated). Highly variable regions were predominantly hypermethylated (**Figure 3B**).

**Figure 3.**
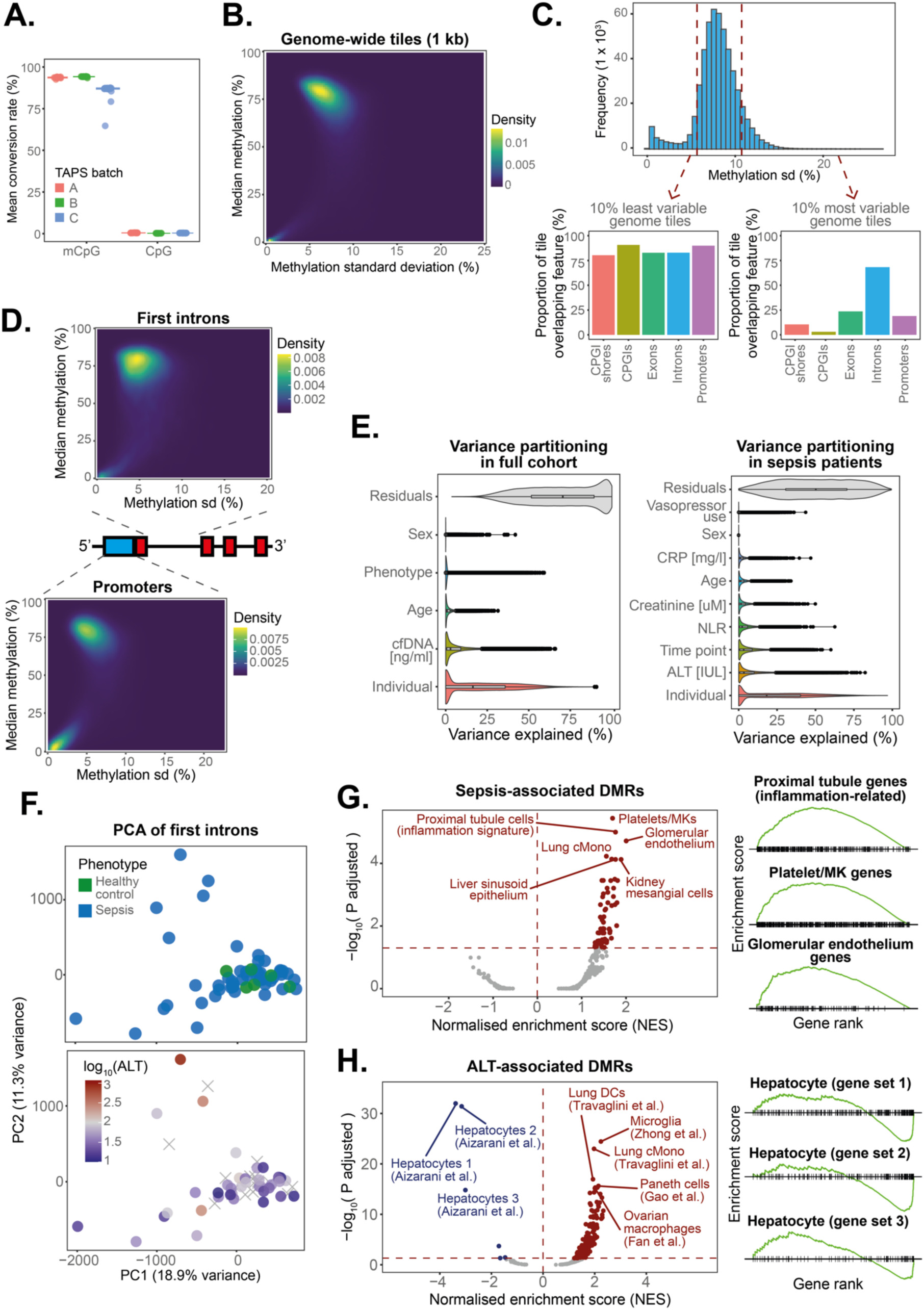
Variation in the cfDNA methylome reflects organ function and disease processes. **A.** mCpG conversion rates in positive and negative spike-in controls as estimated based on our full cohort. Each dot represents a sample, with colours indicating library preparation batch. **B.** 2D-density plot showing the variability (standard deviation, X axis) and average proportion (%, Y axis) of cfDNA methylation at 1-kb genomic windows as estimated based on our full cohort. Shades of colour are proportional to the number of genomic regions located within a given range. **C.** Histogram of cfDNA methylation variability (standard deviation) across all 1-kb windows in the genome (upper panel) as estimated based on our full cohort. The proportional overlap between 1 kb-windows and known genomic annotations is shown for the top 10% most variable and the bottom 10% least variable genomic windows (lower panel). **D.** 2D-density plots showing the variability and average cfDNA methylation at first intron (upper panel) and promoter (lower panel) regions as estimated based on our full cohort. Shades of colour are proportional to the number of regions located within a given range. **E.** The proportion of methylation variance explained by different variables was estimated at intronic regions using variance partitioning analysis. This analysis was conducted separately for the full cohort (left panel) and for sepsis patients only (right panel). Violin plots show the distribution of variance explained by each variable, with dots representing estimates for individual introns. **F.** Principal component analysis based on methylation at first introns. Dots represent samples, with colours indicating either disease status (upper panel) or ALT concentration (lower panel). Crosses indicate samples for which ALT measurements were not available. **G.** Volcano plot showing GSEA-derived enrichment scores (X axis) and FDR-adjusted p values (Y axis) for genomic regions ranked by strength of association between cfDNA methylation and disease status (left panel). Enrichment was estimated based on our full cohort. A subset of gene sets is highlighted. Enrichment score distributions for the topmost enriched gene sets are also shown (right panel). **H.** Volcano plot showing GSEA-derived enrichment scores (X axis) and FDR-adjusted p values (Y axis) for genomic regions ranked by strength of association between cfDNA methylation and ALT levels (left panel). Enrichment was estimated based on sepsis samples only. A subset of gene sets is highlighted. Enrichment score distributions for the topmost enriched gene sets are also shown (right panel).

We next asked if methylation variance was concentrated in any specific types of genomic elements. While lowly variable regions segregated randomly and uniformly across annotations, the top 10% most variable regions were overwhelmingly enriched in introns, with a smaller proportion localising to exons, and promoters (**Figure 3C**). In contrast, variable regions were depleted from CpG islands (CpGIs), which are known to be hypomethylated and invariant (40–43). The high level of variance observed in introns supports recent observations suggesting gene activity correlates most strongly with DNA methylation at the first intron (44). Given that introns accounted for most of the observed variation, we aggregated CpG sites located within the first intronic region of all known genes (**Methods**). In contrast with promoters, which showed evidence of both hyper– and hypo-methylation, first introns showed consistently high methylation proportions and higher levels of variability (**Figure 3D**). We used this intron set as the basis for further exploratory analysis.

We next asked which factors shape the circulating cfDNA methylome during sepsis. We used variance partitioning analysis (45) to identify the variables explaining most of the variability (**Methods**). Surprisingly, only a small proportion of variance (mean = 1.55%) was explained by disease status (sepsis vs controls), with most of the variation being either unexplained or accounted for by inter-individual variability (**Figure 3E**). This could reflect a high contribution of unmeasured factors (e.g. genetic background and infection history) as has been reported for cellular DNA methylation (46,47). It could also relate to disease heterogeneity. We thus repeated this analysis on sepsis patients only, leveraging information from their medical record to quantify the contribution of known clinical factors (**Methods**). While a large proportion of variance remained unexplained, we observed a clear influence of organ function on cfDNA methylation. For example, 1,932 regions showed >30% of their methylation variance explained by circulating levels of alanine aminotransferase (ALT), a marker of liver dysfunction (48–50) (**Figure 3E**). Similar observations were made for creatinine, a marker of kidney dysfunction (51), as well as for C reactive protein (CRP) and the neutrophil-to-lymphocyte ratio (NLR), markers of systemic inflammation (52–54) (**Figure 3E**). This suggests that fluctuations in organ function result in methylation changes in cfDNA at selected genomic regions. Time since ICU admission also had a sizeable impact on methylation variation (**Figure 3E**), suggesting that some methylation changes reflect disease trajectories.

Given its high contribution to methylation variance, we next focused on the impact of liver dysfunction. Principal component analysis of the circulating methylome revealed a clear separation of patients with high ALT measurements from the remaining cohort (**Figure 3F**). In contrast, disease status did not correlate with either principal component (**Figure 3F**). The distribution of samples in PCA space demonstrated a higher level of variability within the patient group than between patients and controls, a phenomenon which could relate to the wide variability in severity of organ dysfunction within sepsis. To test this, we performed differential methylation and gene set enrichment analyses (**Methods**). We identified 4,208 genomic regions where DNA methylation was associated with ALT levels (ALT-associated DMRs) at 5% FDR. Conversely, only 8 regions were differentially methylated between sepsis and controls (sepsis-associated DMRs). Sepsis-associated DMRs were enriched in gene sets related to platelets, epithelium, and inflammation (**Figure 3G**). In contrast, ALT-associated DMRs showed a striking and significant negative enrichment in hepatocyte-specific genes (**Figure 3H**). Since DNA methylation and gene activity are anti-correlated (40,44), this negative enrichment is reflective of a higher hepatocyte contribution to cfDNA. Thus, our observations suggest that patients with high ALT have a significantly higher proportion of hepatocyte-derived DNA in circulation.

Taken together, our observations suggest that the circulating methylome is substantially more variable within sepsis patients than it is between patients and controls, and that a subset of genomic regions show methylation patterns which reflect organ dysfunction and disease trajectories.

### Cell-free DNA accumulation in sepsis is not cell type-specific

We next leveraged methylation profiles to infer the relative contribution of different tissues to circulating cfDNA. We used EpiDISH, a validated methylome deconvolution method (55), to estimate tissue of origin information based on a BS-seq-based reference tissue atlas previously developed for cfDNA deconvolution (31) (**Methods**). Granulocytes showed the highest contribution to cfDNA (37.2% on average), followed by monocytes (11.4%), megakaryocyte-erythroid progenitors (MEPs; 11.3%), circulating lymphocytes, and a much smaller contribution from solid organs (**Figure 4A**). In contrast to other solid tissues, hepatocytes and endothelial cells contributed substantially more DNA to the circulation. These observations agree with previous studies, which have consistently identified erythropoiesis and neutrophil turnover as the two main sources of circulating cfDNA (6,31,56).

**Figure 4.**
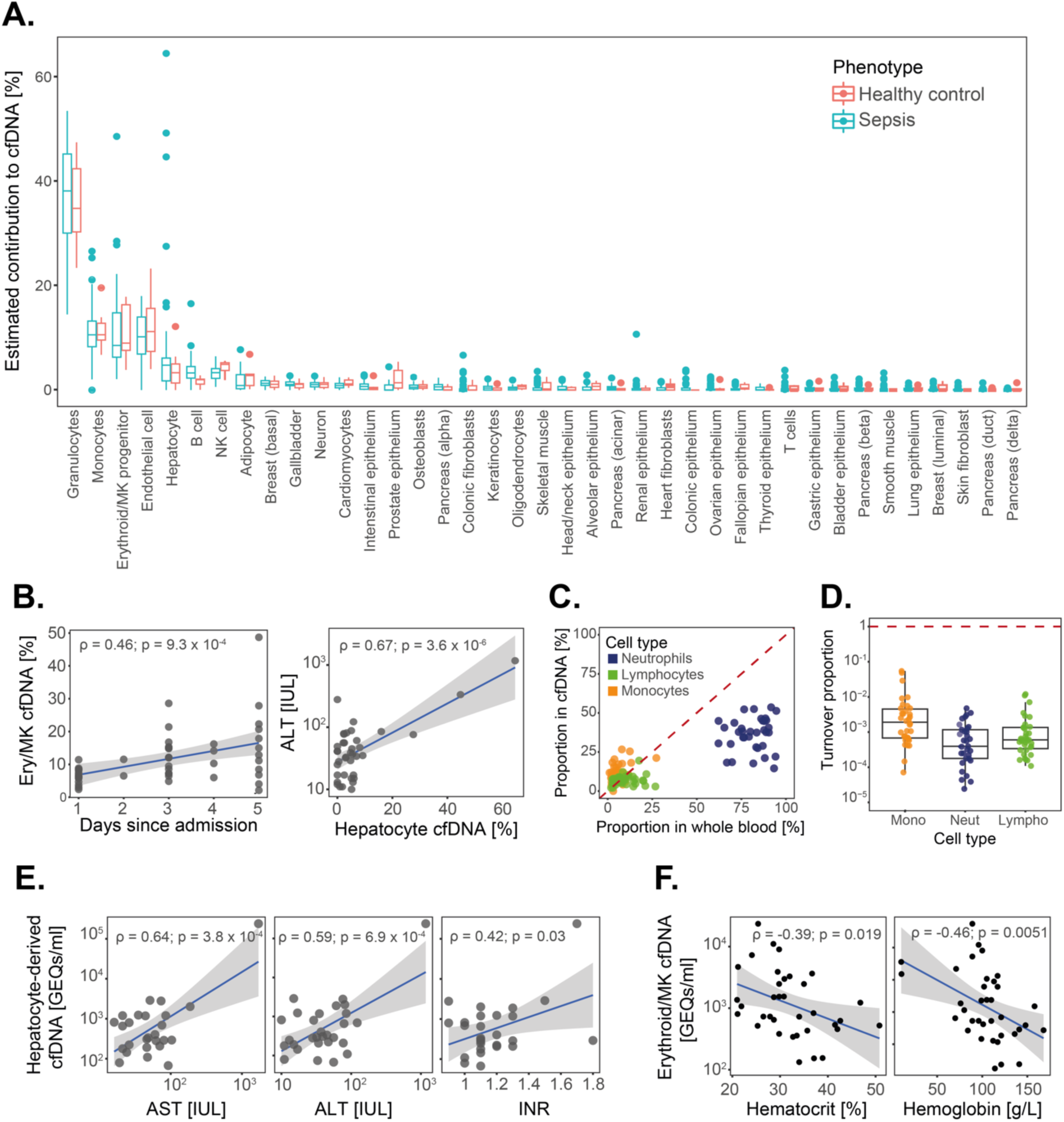
Analysis of tissues of origin of cfDNA during sepsis. **A.** Proportion of cfDNA estimated to arise from different tissues based on methylome deconvolution. Box plots show median and interquartile ranges (IQR) of estimated proportions in sepsis patients (red; n = 49 samples from 31 patients) and healthy controls (blue; n = 7 independent samples). **B.** Correlations between time since hospital admission and proportion of MEP-derived cfDNA (left panel), as well as ALT levels and proportion of hepatocyte-derived cfDNA (right panel). Correlation coefficients and p values were estimated using Pearson correlation tests. **C.** Correlation between proportions of cells in circulation (X axis) and their cellular contributions to cfDNA (Y axis). Colours indicate the three main immune cell lineages. The identity line is shown for reference. **D.** Cellular turnover rates estimated based on cfDNA GEQs and circulating cell proportions. Box plots represent median and IQRs of cellular turnover proportions. The dashed red line indicates a turnover rate of 1, corresponding to cell death and cell generation being equal. **E.** Correlation between markers of liver dysfunction (X axis) and hepatocyte-derived cfDNA GEQs (Y axis). Blue lines and shaded regions linear fits and confidence intervals. Correlation coefficients and p values were estimated using Pearson correlation tests. **F.** Correlation between clinical markers of erythropoiesis (X axis) and MEP-derived cfDNA GEQs (Y axis). Blue lines and shaded regions indicate linear fits and confidence intervals. Correlation coefficients and p values were estimated using Pearson correlation tests.

We next compared cfDNA composition between patients and controls. Surprisingly, we observed no overall difference in cfDNA composition, with cell type contributions ranked in the same order and with similar estimates, regardless of disease status (**Figure 4A**). These results are in stark contrast with previous literature suggesting an increase in granulocyte-derived cfDNA during sepsis (6) and point to accumulation of cfDNA during sepsis not being primarily due to increased immune cell death.

To ensure these results were not biased by technical limitations, we performed further testing to ensure cfDNA composition was robustly and reliably estimated. First, we applied our deconvolution approach to a subset of samples of known cellular composition. To do so, we isolated whole blood leukocytes (WBLs) from 15 sepsis patients and 5 healthy controls and performed DNA methylation profiling (**Methods**). We used these samples to compare cellular proportions estimated from DNA methylation to those directly measured in hospital using a complete blood count (CBC) (**Methods**). In contrast to cfDNA, the whole blood leukocyte methylome was overwhelmingly dominated by cellular composition (**Figure S2A-B**), and estimated cell proportions were highly concordant between methylome deconvolution and direct measurements (**Figure S2C-F**). Moreover, deconvolution successfully distinguished WBL samples from positive controls containing purified neutrophil DNA only (**Figure S2**). These observations greatly strengthened our confidence in our approach.

To further ensure our conclusions were not biased by our choice of deconvolution reference, we repeated our analysis using a different methylation tissue atlas (6) (**Methods**). Results from both atlases were highly concordant, with neutrophils, MEPs, and monocytes remaining the top contributors to the cfDNA pool and cell proportions being largely independent of disease status (**Figure S3**). Thus, we concluded that the observed lack of compositional differences between cfDNA from sepsis patients and controls was unlikely to reflect technical biases and was instead a true biological phenomenon. These observations are incompatible with cfDNA accumulation being a result of higher cell type-specific DNA release (e.g. increased endothelial damage, NETosis or pyroptosis), and instead suggest a potential impairment in cfDNA clearance.

### Variation in cfDNA composition between patients reflects disease processes

We next turned our attention to the relationship between cfDNA and disease features. While overall cfDNA composition did not differ by disease status, we noticed a much larger amount of variation within sepsis patients than in the control group. For example, sepsis patients showed a much broader range of hepatocyte-derived cfDNA proportions, with hepatocytes contributing as much as 60% of DNA in some patients (**Figure 4A**). We observed similarly large variability across cell types, including MEPs, monocytes, and neutrophils. This could reflect disease heterogeneity.

To investigate this, we tested for associations between cfDNA composition and clinical variables (**Methods**). We observed that the proportion of MEP-derived cfDNA was low after hospital admission, but consistently increased over the course of hospitalisation, gradually returning to a value closer to the baseline proportion observed in healthy controls (**Figure 4B**). This could reflect hematopoietic rewiring during the early stages of sepsis, a well-known phenomenon which occurs during infection to support higher production of neutrophils and effector immune cells at the expense of erythrocytes and other hematopoietic lineages (57). Similarly, the proportion of hepatocyte-derived cfDNA was significantly associated with clinical measures of liver dysfunction (**Figure 4B**), suggesting a higher contribution of this cell type in individuals with liver damage. These observations supported our hypothesis that variation in the contribution of different tissues to the cfDNA pool reflects features of disease.

We next integrated our deconvolution results with cfDNA concentration measurements to estimate the number of genomes contributed by each cell type (genome equivalents, GEQs), as has been previously proposed (6) (**Methods**). GEQs are proportional to the number of cell death events occurring within a given cell type, thus providing a principled method to reliably track cellular turnover. We first focused on circulating immune cells. We observed little correlation between the proportion of immune cells in circulation (derived from clinical CBCs) and their proportion in cfDNA (**Figure 4C**). This phenomenon is well documented (58) and supports the theory that cfDNA composition is proportional to cell death, and not to the number of live cells present at any given time. We then estimated cellular turnover rates as the ratio of cell death events (cfDNA GEQs) to live cells (CBC counts) per unit volume of blood (**Methods**). Across all major immune lineages, the number of live cells greatly exceeded the number of cell death events. This was particularly evident for neutrophils, which showed the lowest turnover rates (**Figure 4D**).

These observations agree with the role of emergency granulopoiesis in sepsis, where neutrophil production from hematopoietic stem and progenitor cells (HSPCs) is greatly increased to support the host immune response (57,59).

We then turned our attention to liver dysfunction. We confirmed that hepatocyte-derived cfDNA GEQs were significantly associated with clinical signs of liver dysfunction, including ALT, aspartate aminotransferase (AST), and the international normalised ratio (INR), a measure of prothrombin time, and hence the synthetic function of the liver (**Figure 4E**). Thus, monitoring the levels of hepatocyte-derived cfDNA could be a promising strategy for prompt detection of liver dysfunction. Similarly, we found a significant association between MEP-derived cfDNA GEQs and clinical measures of erythroid production (haematocrit and haemoglobin levels). This implies that measuring MEP-derived cfDNA could enable real-time monitoring of hematopoietic re-wiring and fluctuations in red blood cell (RBC) production.

Taken together, these observations suggest that tracking cfDNA composition is a promising strategy for real-time assessment of disease processes like granulopoiesis, erythropoiesis, and organ dysfunction secondary to sepsis.

### Cell-free DNA fragmentation reveals impaired hepatic clearance during sepsis

Our observations from methylome analysis suggest that cfDNA clearance may be impaired during sepsis. Cell-free DNA is first cleared by Kupffer cells in the liver, a process referred to as hepatic clearance (60). DNA and chromatin fragments released from hepatic clearance are then removed from the bloodstream by the glomerulus (renal filtration), resulting in their excretion as ultra short fragments in the urine (61). Thus, we next asked whether we observed any evidence of hepatic clearance being impaired during sepsis.

Circulating cfDNA exists as small chromatin fragments which originate from apoptosis (62). These fragments, usually one to two nucleosomes in size, are cleaved by extracellular nucleases like DNASE1 and DNASE1L3 upon release into the circulation (63,64). Thus, we reasoned that if hepatic clearance was impaired, cfDNA would remain in circulation for longer, therefore being exposed to cleavage by nucleases for a prolonged period. We hypothesised this would leave a distinct cfDNA fragmentation signature which could be detected using sequencing.

We used read length information to infer the distribution of cfDNA fragment sizes in our patient cohort (**Methods**). In agreement with previous literature (65), most cfDNA fragments (mean=90.75%, median=91.9%, sd=3.25%) were mononucleosomal particles (120-220 bp; mode = 165 bp), with a smaller subset formed of two (mean=5.2%; 220-420 bp; mode = 343 bp) or more (0.61%; > 420 bp; mode = 524 bp) nucleosomes (**Figure 5A**). Mononucleosomal fragments showed evidence of ∼10 bp periodicity, a common feature of cfDNA known to arise from the orientation of DNA grooves around nucleosomes during cleavage (66). This gave us confidence in the quality of our inferred fragmentation information.

**Figure 5.**
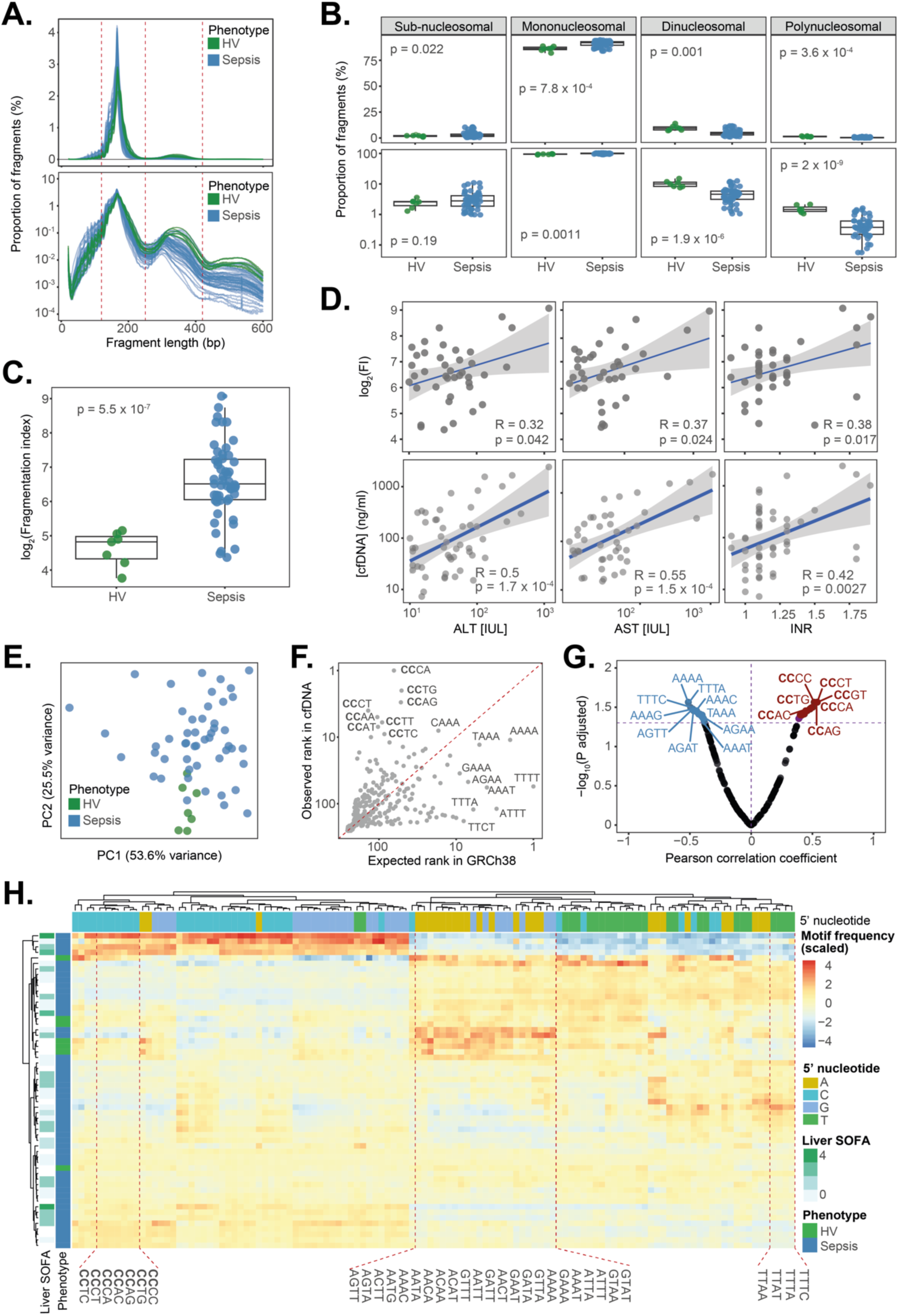
cfDNA fragmentation reveals impaired hepatic clearance during sepsis. **A.** The estimated percentage of cfDNA fragments (Y axis) of different lengths (X axis) in linear (top panel) and logarithmic (bottom panel) scales. Each line represents a sample, with colours indicating control and sepsis patient samples (n = 51 samples from 31 patients and 7 control samples). **B.** Percentage of cfDNA fragments classified as subnucleosomal, mononucleosomal, di-nucleosomal or polynucleosomal based on their observed length. Proportions are shown in both linear (top panel) and logarithmic (bottom panel) scales. Each dot represents a sample, with colours indicating disease status. Box plots show median and IQR values for each sample group. P values were estimated using Wilcoxon Rank-Sum tests. **C.** cfDNA fragmentation indexes (Y axis, in logarithmic scale) stratified by disease status (X axis). Each dot represents a sample, with colours indicating disease status. Box plots show median and IQR values for each sample group. P values were estimated using Wilcoxon Rank-Sum tests. **D.** Correlation between clinical liver function tests (X axis) and either cfDNA fragmentation indexes (Y axis, top panels) or cfDNA concentrations in plasma (Y axis, bottom panels) in sepsis patients. Each dot represents a sample, with blue lines and shaded regions indicating linear fits and confidence intervals. Correlation coefficients and p values were estimated using Pearson correlation tests. **E.** Comparison of average observed (Y axis) and expected (X axis) cfDNA end-motif frequencies in our cohort. Each dot represents a 4 bp 5’ end-motif, with a subset of motifs off the diagonal being highlighted. The identity line is shown for reference. **F.** Principal component analysis plot based on the frequencies of all 4 bp 5’ end motifs. Each dot represents a sample, with colours indicating disease status. **G.** Volcano plot showing the association between end-motif frequency and liver SOFA scores. Each dot represents a motif, with their correlation coefficient with liver SOFA (X axis) and FDR-adjusted p value (Y axis) from Pearson correlation tests shown. Significantly positively and negatively correlated motifs are highlighted in red and blue, respectively. **H.** Heatmap of 5’ cfDNA end-motif frequencies across all samples in our cohort. Shades of colour represent end-motif frequencies, with marginal colour bars indicating the first nucleotide in the motif (horizontal axis), as well as disease status and liver SOFA (vertical axis). Samples were grouped using hierarchical clustering of motif frequencies. A subset of motifs showing appreciable differences in frequency are highlighted.

We next asked if sepsis changed cfDNA fragmentation. Cell-free DNA was significantly more fragmented in sepsis patients, with an enrichment in mononucleosomes and a corresponding depletion in larger fragments (**Figure 5B**). To better describe this skew, we used a previously proposed approach (62,63,67) to calculate cfDNA fragmentation indexes (FIs), defined as the ratio of mononucleosomal to di-nucleosomal fragments (**Methods**). FIs were significantly increased in sepsis (**Figure 5C**), consistent with cfDNA being more fragmented. To understand if this skew was driven by impaired hepatic clearance, we tested for associations between cfDNA fragmentation and liver dysfunction. FIs were significantly correlated with ALT, AST, and INR, as would be expected upon impaired hepatic clearance (**Figure 5D**). This was also evident from the extent of cfDNA accumulation, with cfDNA concentration in plasma being significantly correlated with all measured markers of liver dysfunction (**Figure 5D**). Thus, fragmentation patterns support the hypothesis of impaired hepatic clearance.

Different nucleases are known to show different biases in their cutting site preferences (64). Thus, we hypothesised that impaired hepatic clearance should leave behind a distinctive end-motif signature in cfDNA, reflecting prolonged exposure to specific nucleases. To test this, we leveraged our sequencing data to infer fragment end-motif sequences and to estimate their proportion in cfDNA (**Methods**). In agreement with previous studies, cfDNA end-motifs were significantly enriched in 5’-CCNN sequences, known to be preferentially generated by DNASE1L3, compared to their expected frequency based on the composition of the human genome (**Figure 5F**). This agrees with the known role of DNASE1L3 in cfDNA generation (62,63,67).

We next assessed the impact of sepsis on the end-motif landscape. Principal component analysis of end-motif frequencies showed a clear separation between sepsis patients and controls (**Figure 5E**), indicative of sizeable differences in motif usage. To understand whether the observed differences in end-motif frequencies were driven by impaired hepatic clearance, we tested for associations between end-motif usage and liver-specific sequential organ failure assessment (SOFA) scores, a widely used clinical measure of liver dysfunction (68) (**Methods**). Liver SOFA scores were significantly and positively correlated with known DNASE1L3-associated motifs (i.e. 5’CCNN), supporting the hypothesis that liver dysfunction prevents hepatic DNA clearance (**Figure 5G**), thus exposing cfDNA to prolonged cleavage by DNASE1L3 in circulation. Further exploratory analysis of the end-motif landscape confirmed that 5’CCNN motif frequencies are present at lower frequencies in the control group, increasing during acute sepsis in a way which is proportional to liver dysfunction (**Figure 5H**). Conversely, the proportion of 5’ANNN motifs was significantly higher in the control group (**Figure 5H**). Fragments with 5’-A motifs at both fragment ends have previously been associated with DFFB, a strictly intracellular nuclease known to play a key role during apoptosis (62,63). Thus, our observations are compatible with cfDNA being originally derived from apoptosis, but with impaired clearance during sepsis subsequently diluting this signature from cfDNA due to prolonged exposure to circulating nucleases.

Taken together, our results from fragmentation analysis support the hypothesis that cfDNA accumulates in the circulation during sepsis due to impaired hepatic clearance. This results in cfDNA exposure to circulating nucleases being prolonged, which increases its fragmentation and leaves behind a distinctive end-motif signature. These results highlight gaps in previous literature and illustrate the importance of better understanding cfDNA biology to more accurately interpret fluctuations in the levels of this biomarker.

### Cell-free DNA retains cell type-specific nucleosome positioning signatures

Recent studies have demonstrated that cfDNA retains nucleosome footprints reflective of chromatin architecture in its tissues of origin and have suggested this can be leveraged to infer cfDNA composition (69–72). This holds great promise for cfDNA-based biomarkers, as methylation profiling requires laborious experimental protocols that cannot be easily adapted to clinical settings. In contrast, nucleosome footprinting can be inferred using more widely available DNA-sequencing techniques. This motivated us to map nucleosome footprints, assess whether they retain tissue-specificity, and compare them to our observations from methylome deconvolution.

We adapted an approach by Snyder et al. (69) to estimate windowed protection scores (WPS) around the transcriptional start site (TSS) of all known genes (**Methods**). WPS values are designed to capture the degree of DNA protection from cleavage by nucleases. Because nucleases predominantly cut DNA at the linker region between nucleosomes, WPS reflect nucleosome positioning (69). Analysis of WPS values revealed a clear nucleosome footprint at gene regions (**Figure 6A**). This footprint was characterised by a dip in signal at the TSS, known to be a nucleosome-free region (NFR), and subsequent periodic oscillations which peaked approximately every 180 bp, consistent with the known length of DNA wrapped around each nucleosome (**Figure 6A**).

**Figure 6.**
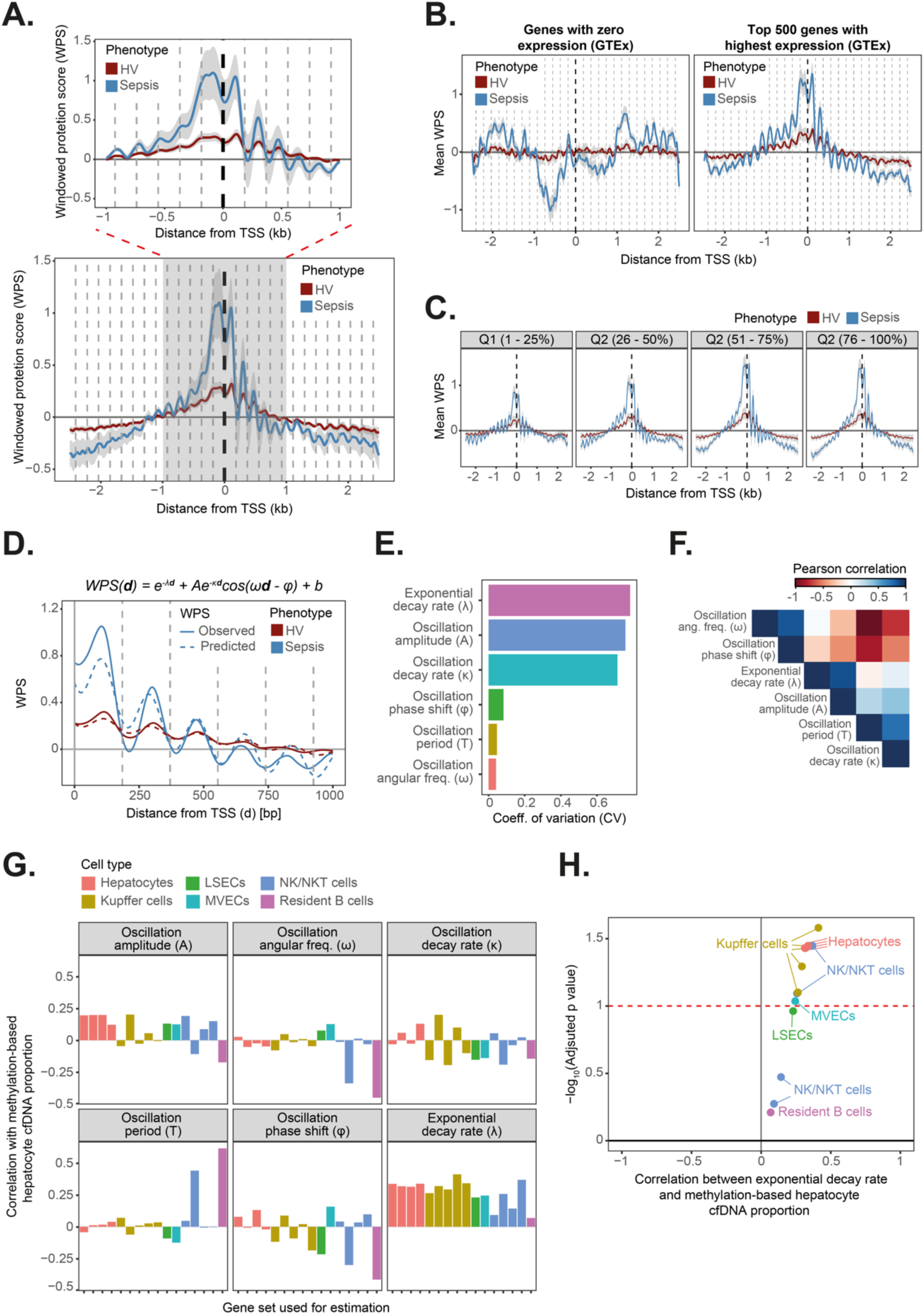
Cell-free DNA retains cell type-specific nucleosome positioning signatures. **A.** Average WPS values and their associated 95% confidence intervals were estimated for all known gene regions using a 5 kb window centred at the TSS. Blue and red lines indicate estimates for sepsis patients and healthy controls, respectively (n = 51 samples from 31 patients and 7 control samples). Dotted vertical lines indicate the expected position of each nucleosome and are spaced 187 bp from each other. A regional plot focusing on a 2 kb region around the TSS is also shown (top panel). **B.** Average WPS values at the TSS regions of active (right panel) and inactive genes (left panel), based on gene expression measurements from the GTEx study. Colours indicate disease status, and dotted vertical lines indicate the expected position of each nucleosome. **C.** Average WPS values at the TSS regions of increasingly more active genes, based on gene expression measurements from the GAinS study. Colours indicate disease status. **D.** Observed WPS values at the TSS regions of all known genes (solid lines) are shown alongside predictions derived from a dampened harmonic oscillator model (dotted lines). Colours indicate disease status. The equation used for model fitting is shown for reference. **E.** Dampened harmonic oscillator models were fitted separately to each sample in our study. The coefficient of variation of estimates for each model parameter are shown as a bar plot. **F.** Heatmap of correlation estimates computed between all model parameter estimates. Colours indicate Pearson correlation coefficients. Rows and columns were ordered by similarity using hierarchical clustering. **G.** Dampened harmonic oscillator models were fitted to the TSS region of tissue-specific gene sets. Bar plots show the Pearson correlation coefficient (Y axis) between each model parameter estimate and the proportion of hepatocyte-derived cfDNA as estimated based on DNA methylation. Each plot shows estimates for a different model parameter, with colours indicating the gene set used during model fitting. **H.** Volcano plot of correlations between hepatocyte-derived cfDNA proportions estimated from methylome deconvolution and WPS exponential decay rates estimated for different liver cell types. The dotted line indicates the statistical significance threshold of BH-adjusted p value < 0.05. Colours indicate the gene set used during model fitting.

Nucleosome positioning at active genes is characterised by the first nucleosome downstream of the TSS (+1 nucleosome) being consistently placed at approximately the same position across cells, a phenomenon known as phasing (73–75). The strength of nucleosome phasing decreases for nucleosomes positioned further away from the TSS, which eventually return to a random arrangement. We confirmed that these features were well captured in our data, with WPS values and oscillation amplitudes weakening proportionally to their distance from the TSS (**Figure 6A**).

Having confirmed that our data retains nucleosome footprints, we turned our attention to the impact of disease status. The strength of nucleosome phasing was substantially higher in sepsis patients compared to controls (**Figure 6A**). We hypothesised that this could be driven by higher levels of gene activity in sepsis, where immune cells and tissues are known to activate a variety of gene expression programs to control infection (15,35,76,77). To confirm that nucleosome footprints were indeed reflective of gene activity, we stratified genes based on their expression level. We used data from the genotype tissue expression (GTEx) consortium to define sets of inactive and active genes (**Methods**). Active genes showed a distinct periodic WPS signal which was sharply increased in sepsis (**Figure 6B**). In contrast, inactive genes showed no difference in WPS between patients and controls, and contained no evidence of nucleosome phasing, as demonstrated by the absence of periodic oscillations or any relationship between WPS and genomic distance (**Figure 6B**). To better understand this, we asked if genes expressed at different levels showed appreciable differences in their nucleosome footprints. We used data from the UK Genomic Advances in Sepsis (GAinS) study (15,20,76) to classify genes into four groups based on their level of expression in whole blood during sepsis (**Methods**). Nucleosome footprint strength progressively increased as we moved from genes in the bottom quartile to genes in the top gene expression quartiles (**Figure 6C**), demonstrating a positive relationship between gene activity and nucleosome phasing.

Taken together, we confirmed that nucleosome footprints are well captured by cfDNA, and that they are indicative of gene activity. Further, we observed evidence of a widespread increase in active gene WPS during sepsis.

Having confirmed our ability to infer gene activity from nucleosome footprints, we sought to develop a principled analytical method to quantitatively study nucleosome phasing using cfDNA data. Previous studies have approached this problem using methods like Fourier series analysis, which can approximate the WPS signal using a combination of trigonometric functions (69,71). However, results from these approaches are often difficult to interpret. Instead, we reasoned that nucleosome positioning could be modelled by combining an exponential decay function with a dampened harmonic oscillator, a model widely used in classical physics (78). In brief, we assume that nucleosome phasing strength decays over genomic distance, that the nucleosome footprint signal oscillates periodically, and that the amplitude of these oscillations decay proportionally to their distance from the TSS (**Methods**). We used non-linear least squares analysis to fit this model to WPS values observed across our cohort (**Methods**). Comparison of observed WPS to the predictions from this approach confirmed that this model adequately captures the periodic oscillations in WPS, as well as the decay in phasing over distance (**Figure 6D**).

To better understand nucleosome phasing, we next fitted models separately to each sample in our study (**Methods**). We confirmed that model fitting performed reliably across the sample set, with high concordance between observed and predicted WPS values (**Figure S4**). We then asked to what extent different nucleosome phasing parameters tended to vary across samples. Parameters describing the oscillation frequency (i.e. phase shift, period, and angular frequency) showed little variability, as indicated by their coefficients of variation (CVs) being close to 0. This reflects the fact that these parameters relate to nucleosome size, which is fixed and invariant (**Figure 6E**). Reassuringly, the median oscillation period inferred by our models was 182 bp (median=182 bp, mean=183.1 bp, sd=8.52 bp), which matches the known length of DNA wrapped around each nucleosome. In contrast, parameters describing nucleosome phasing (i.e. oscillation amplitude and exponential decay rate), as well as the decay in phasing over genomic distance (i.e. oscillation decay rate) showed much higher levels of variability, with CVs > 0.6 (**Figure 6E**).

We next asked if any of these parameters captured important aspects of cfDNA biology, and whether they supported our observations from cfDNA methylation. Visual inspection of the cross-correlation between all model parameters revealed three parameter groups: 1) those related to nucleosome size and spacing (angular frequency and phase shift), 2) those related to signal strength at the TSS (oscillation amplitude and exponential decay rate), and 3) those related to nucleosome phasing downstream of the TSS (oscillation decay rate) (**Figure 6F**). In contrast with the others, parameters in the second group were consistently and significantly increased in sepsis (**Figure S5**), concordant with a generalized increase in gene activity. Thus, we reasoned that parameters describing WPS signal strength at the TSS are the most likely to retain tissue-specific information.

To test this, we repeated our analysis using tissue-specific gene sets derived from previous single-cell studies (**Methods**). Given the importance of liver dysfunction in sepsis, as well as the evidence of hepatic clearance impairment, we focused on liver-specific cell types. We collated gene sets specific to liver cell types from Aizarani et al.’s single-cell liver atlas (79). These gene sets comprised both core structural liver cell types (i.e. hepatocytes, liver sinusoidal endothelial cells (LSECs), microvascular endothelial cells (MVECs)), as well as transient and resident immune cells (Kupffer cells, resident B cells, NK and NKT cells). We fitted separate models to each of these gene sets and assessed the correlation of the inferred parameters with methylome-derived hepatocyte proportions (**Methods**). This enabled us to identify the parameters that best capture tissue-specific gene activity, as well as to assess the agreement between conclusions derived from DNA methylation and nucleosome positioning. While most parameters showed little evidence of correlation, we observed consistently positive and high correlation estimates between methylation-derived hepatocyte proportions and nucleosome phasing exponential decay rates across most tested cell types (**Figure 6G**). This confirmed our hypothesis that signal strength at the TSS is reflective of tissue-specific gene activity, and that this parameter correlates well with estimates derived from DNA methylation. Further inspection revealed very strong statistical evidence of correlation for core liver cell types (hepatocytes, MVECs, LSECs), but non-significant correlations across most immune cells (resident B cells and NKT cells). Kupffer cells, the resident liver macrophages in charge of degrading cfDNA, were a clear exception, showing the highest correlation estimates (**Figure 6H**).

To further confirm this, we visually inspected nucleosome footprints for each of these gene sets, stratifying samples into three groups based on their levels of ALT in circulation. Observations from this analysis confirmed that higher ALT is accompanied by a more pronounced nucleosomal footprint in core liver cell types and Kupffer cells, but not in other liver immune cells (**Figure S6**). These observations demonstrate that nucleosome positioning retains tissue-specific information which can be leveraged to estimate the tissues of origin of cfDNA. Moreover, they suggest that liver damage secondary to sepsis results in an increase in circulating cfDNA derived from the liver parenchyma and from Kupffer cells. Importantly, this increase in Kupffer cell death could explain why hepatic clearance of cfDNA is impaired in these patients.

Taken together, we showed that cfDNA data retains nucleosome footprints, and that these contain tissue-specific information reflective of cfDNA’s cellular context of origin. We developed a principled approach to quantitatively study these footprints and proposed that parameters which relate to footprint strength at the TSS of cell type-specific genes can be used to reliably estimate the contribution of these cell types to circulating cfDNA.

## Discussion

Circulating cfDNA is a promising biomarker for precision medicine, as demonstrated by its widespread use in cancer, pregnancy, and transplantation (2,2,4,5,9,10). However, despite studies showing its relevance in acute infection and critical illness (21,23,80–82), the clinical utility of cfDNA in sepsis has not been explored. Here, we present the first high-throughput multi-modal study of cfDNA during acute sepsis, comprising an in-depth analysis of circulating methylomes, fragmentation landscapes, and nucleosome footprints in a cohort of 56 samples from 31 hospitalised sepsis patients and 7 healthy controls. These data allowed us to delineate the sepsis circulome and provides a resource and point of reference for future studies of circulating nucleic acids in critical illness.

Our observations demonstrate a striking accumulation of cfDNA in the circulation during sepsis and show that the levels of cfDNA in plasma reflect severity of illness. Moreover, we successfully leveraged DNA methylation and computational deconvolution methods (33,83) to infer the relative contribution of different tissues to the cfDNA pool and to study cellular turnover rates (58), demonstrating that cfDNA composition reflects disease processes such as hematopoietic re-wiring to support increased neutrophil production and reduced erythropoiesis during the early stages of disease, as well as instances of organ dysfunction such as liver damage. Surprisingly, and in contrast with previous studies (6), we found no evidence of the observed cfDNA accumulation being driven by increased cell type-specific death such as NETosis, pyroptosis, or endothelial damage. Instead, we found cfDNA composition to be similar between patients and controls. Given the larger sample size of our study, as well as the fact that previous studies had to combine samples from multiple patients to achieve enough coverage (6), we believe our observations greatly expand on previously published data and clarify potentially confounding observations. We hypothesise that such an increase in cfDNA levels in the absence of compositional differences could only be explained by impaired hepatic clearance of DNA and nucleosomal particles from the blood during sepsis. This hypothesis is supported by cfDNA fragmentation and end-motif patterns, both of which show evidence of prolonged exposure of cfDNA to cleavage by circulating nucleases in a way which is directly proportional to all measured markers of liver dysfunction. While the discovery of such a prominent role for liver-mediated clearance in shaping the cfDNA landscape is surprising, it is in line with recent *in vivo* studies, which show that administering therapeutic agents that block hepatic clearance can substantially increase the amount of DNA recovered from liquid biopsies (84). This therapeutic blockade of cfDNA clearance is reminiscent of our observations in sepsis.

In addition, we demonstrated that cfDNA data retains nucleosome footprints, and that these in turn contain information regarding cell type-specific gene activity. While this has been independently demonstrated by previous studies (69,71), here we present a principled approach to study nucleosome phasing and infer cell type-specific information which relies on an exact mathematical model instead of on numeric approximations. When integrating our approach with cell type specific gene sets derived from single-cell studies, we observed that sepsis patients with clinical evidence of liver dysfunction have a higher contribution of DNA from the liver parenchyma and Kupffer cells to the circulation. We believe this can serve as a proof of concept for the development of more complete nucleosome-based cell type deconvolution approaches.

While our study greatly expands our understanding of how the cfDNA landscape fluctuates during sepsis, it is important to acknowledge its limitations. Firstly, our sample size is still limited, and thus we are constrained by statistical power to only detect large changes in the circulome. Future studies with larger sample sizes will be instrumental to study less frequent forms of organ failure and develop statistically validated diagnostic tools. Secondly, while we provide substantial evidence of an impairment in hepatic clearance of cfDNA during sepsis, it is at present unclear which mechanisms are responsible for this phenomenon. These could include increased Kupffer cell death secondary to sepsis; Kupffer cells being co-opted to fight the underlying infection at the expense of reduced efferocytosis; reduced blood flow into the liver due to cardiovascular alterations and shock; and occlusion of hepatic blood flow by immune complexes or NETs. Functional studies dissecting Kupffer cell function and imaging studies assessing liver perfusion are needed to clarify this. Thirdly, it is at present unknown how much of the cfDNA landscape is genetically controlled. In the future, population-wide studies will be needed to map the genetic architecture of cfDNA features such as fragmentation, end-motif frequency, and DNA methylation. Fourthly, despite its advantages, TAPS remains too lengthy a protocol for use in clinical settings. New long-read technologies which enable direct methylation profiling with rapid sample preparation protocols (e.g. Oxford Nanopore and Pacific Biosciences sequencing) could help achieve reliable cfDNA profiling with much shorter turnaround times (56,85–90). Moreover, long-read technologies can also be used to detect less frequent epigenetic marks such as 5hmC and 6mA, which are known to be more cell type-specific than 5mC (91–93). Finally, pathogen-derived DNA is also present in circulation (94,95). Profiling this material for metagenomic analysis could greatly advance diagnostic accuracy and fine-tune antibiotic usage (96). While this can be explored using existing data, current protocols are not best placed to profile this material, which is often highly damaged. Techniques specifically developed for damaged DNA profiling (e.g. single-stranded library preparation methods) could help bridge this gap (97).

## Acknowledgments

We thank all the patients, patient families, nurses, and clinicians who participated in the Sepsis Immunomics and GAinS studies.

## Author contributions

Conceptualization: KCG, JCK

Supervision: JCK, CXS

Data analysis: KCG, PM, EF

Interpretation of results: KCG, PM, MI, JCK

Experimental work: KCG, PM, MI, SH, EF, CW

Patient recruitment: JCK, HQ, SM

Writing: KCG, PM, JCK

## Software, data and materials availability

All codes used for processing and analysis of cfDNA TAPS data are publicly available on GitHub (https://github.com/jknightlab/TAPS-pipeline/).

TAPS-derived cfDNA methylation matrices containing estimated methylation proportions at 19,288,064 autosomal and variable CpG sites, as well as their associated metadata, are publicly available on Zenodo (https://doi.org/10.5281/zenodo.14844792). Raw sequencing files in FASTQ format for the data presented in this study will be deposited in the European Genome-Phenome Archive (EGA) upon study publication.

## Methods

### Study design and participants

#### Sepsis Immunomics cohort

The Sepsis Immunomics (SI) study is an observational study set up in Oxford to better understand the host response during sepsis via the use of immunological and high throughput omic approaches (South Central Oxford REC C, reference:19/SC/0296). Patients were eligible to join the SI study if they were ≥ 18 years of age and admitted to hospital with suspected sepsis at Oxford University Hospitals NHS Foundation Trust, UK. Recruitment was conducted at the intensive care unit (ICU), emergency department (ED), and hospital wards. In the ICU, patients were recruited if they had symptoms and signs of established sepsis (i.e. suspected infection and an acute change in SOFA score ≥ 2). In the ED and medical wards, patients were deemed eligible if they had suspected infection and a change in quick SOFA score (qSOFA) ≥ 2, as well as a National Early Warning System (NEWS2) score ≥ 7. Patients were excluded if consent could not be obtained; if there was an advanced directive to withhold life-sustaining treatment; when admitted to hospital for palliative care only; if pregnant or within 6 weeks post-partum; if presenting with immunodeficiency due to steroid therapy, HIV infection or other immunosuppressive agents; and if diagnosed with metastatic disease or haematological malignancy.

Acute samples were collected at admission and repeatedly at regular intervals within the first 9 days of hospitalisation, most often at days 3 and 5 post-admission. Patients contributing data to the analyses in this publication were recruited between 2022 and 2024.

#### Healthy volunteer cohort

Volunteers self-reporting as healthy (HV) and with no recent history of infection were recruited at the Centre for Human Genetics, University of Oxford, following informed consent (ethical approval South Central Oxford REC B, reference 06/Q1605/55).

### Sample collection and processing

#### Collection of plasma

Blood samples were collected in EDTA vacutainer tubes (BD Biosciences) and processed within 1 hour of collection. For plasma isolation, 10 ml of blood were centrifuged at 1,600 g for 10 minutes at 4°C. Plasma was obtained from the top layer following centrifugation and clarified by performing a second, high-speed centrifugation at 16,000 g for 10 minutes at 4°C to remove remaining cellular debris. Plasma was then aliquoted into 1.5 ml microtubes and stored at –80°C until DNA isolation.

#### Collection of whole blood leukocytes and CD66+ cells

In a subset of individuals, whole blood leukocytes (WBLs) were obtained from EDTA-stabilised blood by separating the buffy coat layer after centrifugation at 1,600g for 10 minutes. In some individuals, neutrophil enrichment was also performed using the EasySep Whole Blood CD66b+ Positive Selection Kit (StemCell Technologies) according to the manufacturer’s instructions. WBLs and CD66b+ cells were stored at –80°C until DNA isolation.

#### Cell-free DNA isolation from plasma

Circulating cell-free DNA (cfDNA) was isolated from 1 to 3 ml of high-grade plasma, as determined by material availability for any given patient. Isolation was performed using the QIAamp Circulating Nucleic Acid kit (Qiagen) according to the manufacturer’s instructions. Carrier RNA was not added to avoid reads being taken up by the additional RNA during sequencing. DNA was eluted in 35 µl of elution buffer (Buffer EB; Qiagen) and stored at –20°C until library preparation.

Cell-free DNA concentration was estimated fluorometrically using the Qubit High Sensitivity dsDNA assay (ThermoFisher Scientific). DNA quality was assessed using capillary electrophoresis with the TapeStation 4200 system (Agilent Technologies) in combination with the cell-free DNA ScreenTape kit (Agilent Technologies). This enabled estimation of cfDNA purity based on the distribution of fragment sizes. Only samples with cfDNA purity ≥ 75% were taken forward for library preparation.

#### DNA isolation from whole blood leukocytes

Whole blood leukocyte genomic DNA (gDNA) was extracted from 100 µl of frozen buffy coat or CD66b+ enriched cells using the NEB Monarch Genomic DNA Purification kit (New England Biolabs) according to the manufacturer’s instructions. DNA concentration was estimated using the Qubit High Sensitivity dsDNA assay (ThermoFisher Scientific), and DNA quality was inspected with a TapeStation 4200 system (Agilent Technologies) and the genomic DNA ScreenTape kit (Agilent Technologies).

#### Preparation of TAPS spike-in controls

Carrier DNA for TAPS library preparation was derived from PCR amplification of the pNIC28Bsa4 plasmid (Addgene) (reaction composition: 1 ng template DNA, 0.5 µM forward and reverse primers, 1X Phusion High-Fidelity PCR Master Mix (ThermoFisher Scientific) in HF buffer).

Positive and negative spike-in controls (CpG-methylated lambda DNA and 2-kb unmethylated DNA, respectively) were fragmented in a Covaris M220 ultrasonicator (Covaris, LLC). Size selection was then performed using a 1.2X ratio of AmpureXP beads (Beckmann Coulter), with the final median DNA fragment being 200 bp.

#### TAPS library preparation

Between 10 and 50 ng or cfDNA were spiked-in with 0.15% CpG-methylated lambda DNA and 0.015% unmodified 2-kb control. DNA was next end-repaired and A-tailed using the NEBNext Ultra II End Repair/dA-Tailing Module (New England Biolabs). Ligation to Illumina Multiplexing adapters was then performed using the KAPA HyperPrep kit (Roche Sequencing Solutions) according to the manufacturer’s instructions. Subsequently, ligated libraries were subjected to two rounds of oxidiation with mTet1CD and reduction with pyridine borane (PyBr). In brief, DNA was oxidized by incubating it with mTet1CD in a 50 µl reaction for 80 minutes at 37°C (reaction composition: 50 mM Hepes buffer, 100 µM ammonium iron sulfate, 1 mM α-ketoglutarate, 2 mM ascorbic acid, 2 mM dithiothreitol,

100 mM NaCl, 1.2 mM adenosine triphosphate, and 4 µ Tet1CD). Next, 0.8 IU of proteinase K (New England Biolabs) were added, and the reaction was incubated for 1 hour at 50°C. Reaction products were cleaned-up using a Bio-spin P-30 Gel Column (BioRad) and a 1.8X ratio of AMPure XP beads (Beckmann Coulter). Finally, Oxidized DNA was reduced by incubating it with PyBr in a 50 µl reaction for 16 hours at 37°C and 850 rpm agitation (reaction composition: 35 µl nuclease-free water, 600 mM sodium acetate solution, 1 M PyBr). Reaction products were purified using Zymo-Spin columns (ZymoResearch).

The final reaction products were PCR-amplified using the KAPA HiFi HotStart Uracil+ ReadyMix PCR kit (Roche Sequencing Solutions) using a pair of custom-made IDT primers (FW: 5’-AATGATACGGCGACCACCGAGATCTACAC-3’, RV: 5’-CAAGCAGAAGACGGCATACGAGAT-3’). Amplification comprised four PCR cycles, after which DNA was cleaned up using a 1X ratio of AMPure XP beads (Beckmann Coulter).

#### Targeted EM-seq library preparation

Enzymatic methyl-seq (EM-seq) was performed on a subset of samples for which WBL-or neutrophil-derived gDNA was available. To this end, 200 ng of gDNA were sheared using a Covaris M220 ultrasonicator (Covaris, LLC), producing a median fragment size of 200 bp. The NEBNext Enzymatic Methyl-seq Methylation Library Preparation Kit (New England Biolabs) was then used for library preparation according to the manufacturer’s instruction. In brief, DNA was end-repaired, ligated to sequencing adapters, and multiplexed using combinatorial indexing. Indexed DNA was next supplemented with positive and negative spike-in controls (fully methylated *E. coli* pUC19A DNA and unmethylated lambda phage DNA, respectively), and enzymatic methylation was performed on this DNA using a two-step protocol consisting of oxidation of methylated cytosines with TET2, followed by deamination of unprotected (i.e. unmethylated) cytosines with APOBEC. This resulted in methylated cytosines retaining their identity and unmethylated cytosines being converted to uracil and sequenced as thymidines.

Converted DNA was amplified for five PCR cycles (reaction conditions: initial denaturation at 98°C for 30 seconds; five cycles of 1) denaturation at 98°C for 10 seconds, annealing at 62°C for 30 seconds, 2) extension at 65°C for 60 seconds; and final extension at 65°C for 5 minutes).

Amplified DNA was then hybridised to a panel of probes for targeted enrichment using the Twist Target Enrichment Fast Hybridisation protocol (Twist Biosciences). Individual libraries were combined into 8-plex pools for hybridisation with streptavidin baits comprising the Twist Human Methylome panel (Twist Biosciences), covering approximately 4 million CpG sites spread over 123 Mb of the genome.

Following hybridisation, library pools were magnetically purified and DNA was further amplified using 8 cycles of PCR (reaction conditions: initial denaturation at 98°C for 45 seconds; eight cycles of 1) denaturation at 98°C for 15 seconds, annealing at 60°C for 30 seconds, 2) extension at 72°C for 60 seconds; and final extension at 72°C for 1 minute). Final library pools were adjusted to the same DNA concentration and taken forward for sequencing.

#### DNA sequencing

TAPS (cfDNA) and EM-seq (WBL) DNA libraries were sequenced using a NovaSeq X (Illumina) and NovaSeq 6000 (Illumina) instrument, respectively. In both instances, 150 bp paired-end reads were used.

### TAPS data analysis

#### Read alignment and filtering

Sequencing reads were first merged into a single file (FASTQ) per sample. Sequencing adapter trimming was next performed with TrimGalore (v0.6.10) using default parameters. Reads were aligned to a reference containing the human genome sequence (GRCh38, NCBI) and the spike-in sequences used during TAPS library preparation. Read alignment was performed using the Burrows-Wheeler algorithm (98) as implemented in BWA-MEM (v0.7.17). Aligned reads were filtered by mapping quality (MAPQ score ≥ 10) and sorted by genomic coordinate using SAMtools (v1.8) (99). Finally, PCR and optical duplicates were marked and discarded from further analyses using Picard’s (v2.23) *MarkDuplicates* function.

Following alignment, mapped reads were used to identify potential sample swaps using Picard’s (v2.23) *CrosscheckFingerprints* function.

#### Methylation calling

Sequencing library construction comprises an end-repair step which fills in 5’ overhangs and removes 3’ overhangs from input DNA. This results in methylation events being missed when they are located within the overhang of a DNA fragment, a phenomenon referred to as methylation bias (*mbias*) (100). We accounted for this by removing any CpG calls derived from read ends. To do so, MethylDackel’s (v0.6.1) *mbias* function was used to assess the distribution of methylation estimates along the position of sequencing reads. This enabled us to define a set of clipping parameters which minimised potential bias originating from the original top (OT) and bottom (OB) strands, respectively (OT R1 start = 5 bp, OT R1 end = 135 bp, OT R2 start = 5 bp, OT R2 end = 115 bp, OB R1 start = 20 bp, OB R1 end = 145 bp, OB R2 start = 35 bp, OB R2 end = 145 bp). MethylDackel (v0.6.1) was used to clip read ends according to these parameters and to call methylation events at all CpG sites detected in clipped reads. Only bases with a Phred score ≥ 5 were kept for this analysis. Methylation calls were outputted in bedGraph and methylKit (101) formats.

Methylation calls were next quality filtered. BEDtools (v2.31) was used to remove any CpG sites mapping to centromeres, gaps in the genome assembly, regions flagged as ‘blacklisted’ by ENCODE (102), or regions with a high density of repeats (based on UCSC Genome Browser’s *repeatMasker* (103)). Because TAPS cannot distinguish methylation events from C>T single nucleotide polymorphisms (SNPs), CpG sites intersecting any common SNPs (i.e. MAF > 1%) reported in the SNP database (dbSNP; v155) were also removed (104).

MethylDackel was designed for analysis of bisulfite sequencing (BS-seq), which relies on conversion of unmethylated cytosines, as opposed to the direct conversion of methylated cytosines performed by TAPS. Thus, methylation calls had to be flipped (i.e. their effect direction reversed). Call flipping was performed using the *awk* command.

Finally, bedGraph files were converted to bigWig format for visualisation in the Integrative Genomics Viewer (IGV) genome browser (105). mehtylKit files were imported into R (v4.3.2) for analysis.

#### CpG quality filtering and summarisation

Genome-wide methylation calls were imported into R (v4.3.2), quality filtered, and pre-processed using methylKit (v1.28.0) (101). First, mCpG>T conversion rates were estimated as the proportion of CpGs detected as methylated in the positive spike-in controls. Only samples with a conversion rate ≥ 80% were kept for analysis. CpG sites were next filtered by visualising sequencing coverage along each chromosome within each sample to identify outlier regions with unusually high coverage. Only CpGs mapping to autosomes and with a coverage of ≤ 25 reads were kept. Read counts per CpG were next normalised for varying coverage, and a union set of CpGs detected in at least 3 samples per group (i.e. sepsis patients or healthy volunteers) was defined using methylkit.

Feature summarisation was performed using methylkit by averaging normalised methylated and unmethylated read counts and estimating methylation proportions across all CpGs located within a given genomic window. Only windows containing ≥ 5 CpGs were kept. Feature summarisation was performed at the level of: 1) 1-kb tiles spanning the whole genome, 2) promoter regions for all annotated genes reported in GENCODE (v43) (106), 3) the first intron of each gene reported in GENCODE (v43) (106), and 4) regions contained in Loyfer et al.’s methylation atlas (31).

#### Methylation exploratory analysis and batch correction

Methylation proportions tiled at the promoter, intron, and 1-kb tile levels were used for exploratory analysis and data visualisation. In brief, standard deviations were estimated per tile across the cohort and tiles were ranked by variability. The overlap between the most/least variable tiles and known functional elements (i.e. CpGIs, CpGI shores, exons, introns, and promoters) was assessed using methylkit and visualised using ggplot2 (v2.3.4.4). Tiles ranked amongst the top 80% by variability were used as a basis for hierarchical clustering and visualisation of sample dendrograms, as well as for dimensionality reduction using principal component analysis (PCA).

Batch effects between library preparation batches were identified based on PCA visualisation, and batch regression was conducted on tiled methylation proportions using the ComBat algorithm (107) as implemented in the SVA package (v3.50). Tiles with methylation proportions estimates > 1 or < 0 after batch correction were manually set to 1 and 0, respectively.

Following batch correction, sample clustering and PCA were recomputed. Principal components were visually explored to dissect their relationship with known clinical and technical covariates using ggplot2.

#### Variance partitioning analysis

Batch-corrected methylation proportions tiled at the promoter and intron levels were used to identify the main sources of variability contributing to the circulating methylome. To do so, the variancePartition (v1.32.5) R package (45) was used to model methylation proportions as a function of known clinical and technical covariates using a linear mixed model. The following equation was assumed during model fitting:

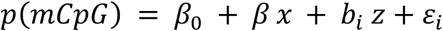

Where β_0_ represents the residual term and ε a random error term, while **x** and **z** stand for variables modelled as fixed and random effects, respectively. When fitting the model across all samples (i.e. both healthy controls and sepsis patients), age and cfDNA yield were modelled as fixed effects, while disease status, self-reported sex, and individual were modelled as random effects. When fitting the model to sepsis samples only, age, time since admission, neutrophil-to-lymphocyte ratio (NLR), peak ALT, peak CRP, and peak Creatinine were modelled as fixed effects, with vasopressor use, self-reported sex, and individual being modelled as random effects.

The estimated proportion of variance explained per variable for each tile was visualised using the variancePartition and ggplot2 packages.

#### Differential methylation and pathway enrichment analyses

Batch-corrected methylation proportions tiled at the promoter and intron levels were used for differential methylation analysis with a moderated T-test, as implemented in limma’s (v3.58.1) *eBayes* function (108). Differentially methylated regions (DMRs) were identified by: 1) contrasting cases and controls, and 2) by modelling methylation proportins as a linear function of ALT values. Regions were deemed differentially methylated if they showed evidence of a statistical association at a false discovery rate (FDR) ≤ 0.05 following multiple testing correction with the Benjamini-Hochberg procedure (109).

The genes corresponding to each promoter or intronic DMR were used as an input for pathway enrichment analysis with the gene set enrichment analysis (GSEA) algorithm (110) using the fGSEA package (v1.28) (111). Enrichment was estimated by comparing the input gene list with a series of cell type-specific gene sets obtained from previous single-cell experiments and maintained by the molecular signatures database (MSigDB) (112). Gene sets were deemed enriched if they showed a non-zero normalised enrichment score (NES) at FDR ≤ 0.05 following multiple testing correction. NES and FDR-adjusted P value estimates were visualised using ggplot2.

#### Deconvolution of cfDNA tissues of origin

We next used methylation profiles to estimate the contribution of different tissues to cfDNA. Methylome deconvolution was performed taking as a reference either: 1) the BS-seq tissue methylation atlas published by Loyfer et al. (31), or 2) a combination of the methylation array-based cell atlases published by Salas et al. (32) and Moss et al. (6).

Deconvolution based on the BS-seq reference was performed by averaging batch-corrected methylation proportions across all CpG sites detected in each region in the atlas. This methylation matrix was used as an input for cell type proportion estimation with the robust partial correlations (RPC) method as implemented in EpiDISH (55) (v2.18.0). Deconvolution based on the array-based references was performed by matching CpG sites to their corresponding Illumina array probe IDs and subsetting the CpG set to those sites present in either Salas’s or Moss’s atlas. The resulting methylation matrix was used as an input for cell type proportion estimation with the RPC method in a hierarchical manner, as implemented in hierarchical EpiDISH (*hepidish*) (83). Because the Moss atlas comprises solid tissues, while the Salas atlas focuses on circulating immune cells, we supplemented Moss’s tissue atlas with an additional column (“*immune cells*”), which was defined as a weighted average of Salas’s atlas, with each cell type being weighted by its average proportion in whole blood. This data set served as a reference for the first deconvolution step. Cell proportions estimated for the “*immune cells*” category were subsequently broken down into cell types by performing a second RPC-based deconvolution step based on Salas’s atlas. This resulted in a final set of cell proportion estimates covering all solid tissues and circulating cells.

Cell-free DNA composition estimates per sample were visualised using ggplot2, and their correlation with clinical and technical variables was assessed in R.

#### Cellular turnover analysis

Results from methylome deconvolution were used to estimate cellular turnover rates. To do so, we estimated the approximate number of genomes contributed by a given cell type per ml of plasma. We refer to this as genome equivalents (GEQs/ml), in line with previous studies (6). Genome equivalents were calculated as follows:

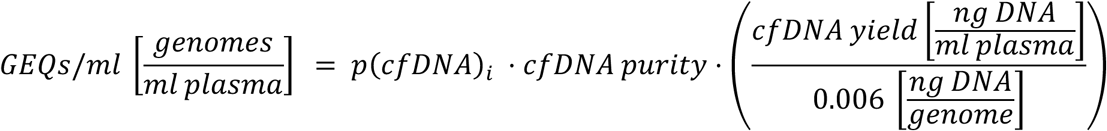

Where **p(cfDNA)**_i_ represents the proportion of cfDNA contributed by cell type **i, cfDNA purity** the proportion of DNA estimated to be true cfDNA based on its fragment size distribution, **cfDNA yield** the amount of DNA obtained per ml of plasma, and **0.006** a constant representing the average DNA contained by a typical cell. GEQs/ml are proportional to the number of cell deaths occurring within a cell type.

Cellular turnover was estimated by integrating cfDNA GEQs/ml with absolute cell counts from clinical blood counts (CBC) carried out for the same samples. Turnover proportions were defined as the ratio of cfDNA GEQs/ml to CBC-based cells/ml for each cell type.

#### Analysis of cfDNA fragmentation patterns

Following alignment and quality filtering, the genomic coordinates of each sequencing read were extracted using SAMtools (v1.8). Read start and end positions were used to infer the length of the corresponding cfDNA fragment. Fragment coordinates were then imported into R and read counts were tabulated against fragment length using tidyverse (v2.0.0), resulting in estimates of the proportion of cfDNA molecules of different lengths. Fragment length distributions were visually inspected using ggplot2.

Fragmentation patterns were contrasted between sepsis cases and controls by comparing the proportion of fragments which fell within subnucleosomal (<120 bp), mononucleosomal (120-250 bp), dinucloeosomal (251-420 bp), or polynucleosomal (>420 bp) size ranges. Proportions were compared between groups using T-tests.

Fragmentation indices (FI) were estimated in line with an approach introduced previously (64). In brief, we defined fragmentation indices as the ratio of the peaks in the fragment size distribution corresponding to mononucleosomal and dinucleosomal fragments:

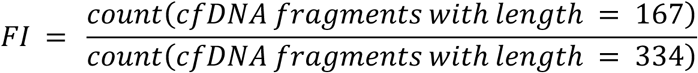

FIs were visualised using ggplot2, compared between groups using T-tests, and tested for associations with continuous clinical variables using Pearson correlation tests.

#### Analysis of cfDNA fragment end-motif frequencies

Fragment end-motifs were obtained following the approach introduced by Jiang et al. (3). In brief, read start and end coordinates were used to fetch the reference genome sequence at the genomic location of interest using UCSC’s *twoBitToFa* function. The first and last 4 nucleotides of this sequence were extracted using pattern matching with *grep*. To account for 5’ overhang filling and 3’ overhang removal during library preparation, end-motifs were converted to the 5’ –> 3’ orientation. To do so, the first four nucleotides (i.e. those mapping to the beginning of the 5’ –> 3’ strand of the reference genome) were taken directly, while the last four nucleotides (i.e. those mapping to the end of the 5’ –> 3’ strand of the reference genome), were reversed in order and their complimentary base pair sequence was derived using character substitution with bash’s *tr* command. This resulted in both fragment end-motifs being in the 5’->3’ orientation, thus representing the true sequence at the end of the fragment overhangs before end repair.

End-motif sequences were imported into R and tabulated, which enabled estimation of their frequency in each sample. To interpret the observed frequencies, BBMap’s exact k-mer frequency calculation algorithm was used to compute expected frequencies for all 4-mer combinations based on the human reference genome sequence. Expected and observed frequencies were compared using scatterplots in R. End-motif frequencies were further used for dimensionality reduction with PCA.

Differential end-motif usage was assessed by contrasting frequencies between sepsis cases and healthy controls using Wilcoxon Rank Sum tests, along with estimation of log_2_-fold changes (LFCs) in frequency for each motif. Wilcoxon p values were corrected for multiple testing using Benjamini-Hochberg’s procedure. Associations between end-motif usage and quantitative clinical variables were identified using Pearson Correlation tests, with correlation p values adjusted for multiple testing using Benjamini-Hochberg’s procedure. Differential end-motif usage was defined as evidence of different end-motif frequencies at an FDR ≤ 0.05.

#### Cell-free DNA-based nucleosome footprinting

Nucleosome footprinting based on cfDNA coverage was performed following the approach by Snyder et al. (69). Briefly, sequencing reads were first filtered based on length to only include information from mononucleosomal fragments (i.e. fragment size ≥ 120 bp and ≤ 200 bp). Next, we tiled gene-containing regions by creating a 120-bp sliding window, which was subsequently slid by 1 bp steps across a 5 kb region centred at the TSS of each gene in GENCODE (v43). The overlap between mononucleosomal fragments and each sliding window was next computed, with fragments fully spanning the window (i.e. with start and end points outside the window) being tabulated separately from non-fully spanning fragments (i.e. fragments with a start or end position within the window). Windowed protection scores (WPS) were estimated as proposed by Snyder et al. (69). Namely, WPS was defined as the number of fully spanning fragments minus the number of non-fully spanning fragments. WPS values were adjusted for sequencing coverage differences by subtracting the mean WPS value for the region, after which they were smoothened using a normally distributed kernel with bandwidth set to 30 bp.

#### Analysis of nucleosome footprints for different gene sets

Cell type-specific and disease-specific gene sets were collated from the following studies: 1) the Genotype-Tissue Expression (GTEx) project (v10) (113), 2) the Genomic Advances in Sepsis (GAinS) study (20,76), and 3) the MSigDB database (112). GTEx median TPMs were used to define a set of ‘*inactive genes’* (i.e. genes with zero expression across all GTEx tissues) and a set of ‘*highly active genes’* (i.e. top 500 genes with largest average TPM values across tissues). Bulk RNA-seq data from the GAinS study were used to group genes into quartiles based on their mean expression level in whole blood of sepsis patients. Finally, MSigDB signatures were subset to cell type-specific gene sets from Aizarani et al.’s single-cell liver atlas (79). Nucleosome footprints were assessed separately for each gene set of interest.

#### Modelling of nucleosome phasing using a harmonic oscillator

We propose that nucleosome phasing downstream the TSS can be captured by the average adjusted WPS value at each position for a given gene set, and that it oscillates periodically as a function of genomic distance. We believe this behaviour is well described by the following model:

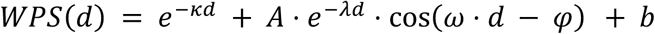

Where **d** represents distance from the TSS (in bp). The first term models the general decay in WPS signal over distance, with **κ** being the exponential decay constant. In contrast, the second term models the periodic oscillation of the WPS signal, with **ω** being the angular frequency of oscillations (in radians) and **φ** the cosine phase displacement (also in radians). The magnitude of the oscillation, **A,** decreases over genomic distance, with this decrease being modelled by a further exponential function with decay constant **λ**. Thus, the second term functions as a dampened harmonic oscillator. Finally, **b** stands for the residual term.

This model was fitted to the observed WPS values in a two-stage process to aid model convergence. Fitting was performed with non-linear least squares (NLS) analysis using R’s *nls* function. The exponential decay term was fitted first, with the dampened harmonic oscillator term being subsequently fitted to the residuals from the first regression. Model fitting was performed separately for each combination of gene sets and samples, with coefficient estimates from each analysis being extracted for future comparisons. Only WPS observations in the region spanning –185 to 925 bp from the TSS were used for model fitting. These capture the full NFR around the TSS, along with the first five downstream nucleosomes.

Model fits were assessed by substituting the estimated coefficients into the model equation and using it to predict WPS as a function of genomic distance. Observed and predicted values were visually compared using ggplot2.

Estimated coefficients were compared between sepsis cases and controls using Wilcoxon Rank Sum tests. Associations between model coefficient estimates and quantitative clinical variables were inspected using Pearson Correlation coefficients. Correlations were subsequently statistically tested using a Pearson Correlation Test, with p values adjusted using Benjamini-Hochberg’s procedure (109).

### EM-seq data analysis

#### Adapter trimming, read alignment, and methylation calling

Raw sequencing reads were processed using nf-core’s methyl-seq pipeline (v2.3.0) (114). This pipeline performs adapter trimming, read mapping, and methylation calling. Briefly, sequencing adapters were trimmed using TrimGalore (v0.6.10) with default parameters. Reads were next aligned to a reference containing the human genome sequence (GRCh38, NCBI) and the spike-in sequences used during EM-seq library preparation. Read alignment was performed using Bismark (v0.22.3) (115), which is optimised for BS-seq data analysis and is thus compatible with EM-seq’s chemistry. Bismark removes duplicate reads, aligns reads to the reference genome, clips read ends to account for methylation bias (see TAPS library preparation methods), and identifies methylation events at C residues covered by clipped reads. Clipping parameters were set to 12 bp from each end of sequencing reads.

Methylation events were called per strand across all cytosine contexts. As CHG and CHH methylation were minimal, only CpG sites were kept for further analysis. Ratios of converted to protected CpGs were subsequently calculated and reported as methylation proportions. Mirrored CpG sites on opposite strands were collapsed into a single analytical unit using Bismark’s *coverage2cytosine* function.

#### CpG quality filtering

Collapsed CpG methylation calls were imported into R using methylKit (v1.28.0) (101). Only CpG sites with a coverage ≥ 10 reads were kept for analysis. Outlying CpG sites in the top 1-percentile by coverage were also removed. Sites were also excluded if they overlapped any common (i.e. MAF > 1%) C>T SNPs reported in dbSNP (v151) (104).

#### Variance partitioning analysis

Methylation proportions were used to identify the main sources of variability contributing to the whole blood methylome using variancePartition (v1.32.5) (45). Methylation proportions were modelled as a function of known clinical and technical covariates using a linear mixed model as detailed above (see TAPS data analysis methods). The following variables were included in the model: age, cellular composition, phenotype (i.e. acute sepsis vs controls), sex, and smoking status. Quantitative variables were modelled as fixed effects, with categorical variables modelled as random effects. Cellular composition was defined as the coordinate of the principal component most associated with CBC-measured blood cell proportions during exploratory analysis of the whole blood methylome.

#### Deconvolution of whole blood leukocyte methylomes

Deconvolution was performed on EM-seq data from WBL and neutrophil samples with the RPC method as implemented in EpiDISH (v2.18.0) (83). The cell type methylation atlas of circulating immune cells published by Salas et al. (32) was used as a reference for this analysis. Cell proportion estimates per sample were visualised using ggplot2, and their correlation with the cell proportions reported in each patient’s complete blood cell counts (CBC) was assessed using Pearson Correlation tests.

## Supplementary materials

**Figure S1.**
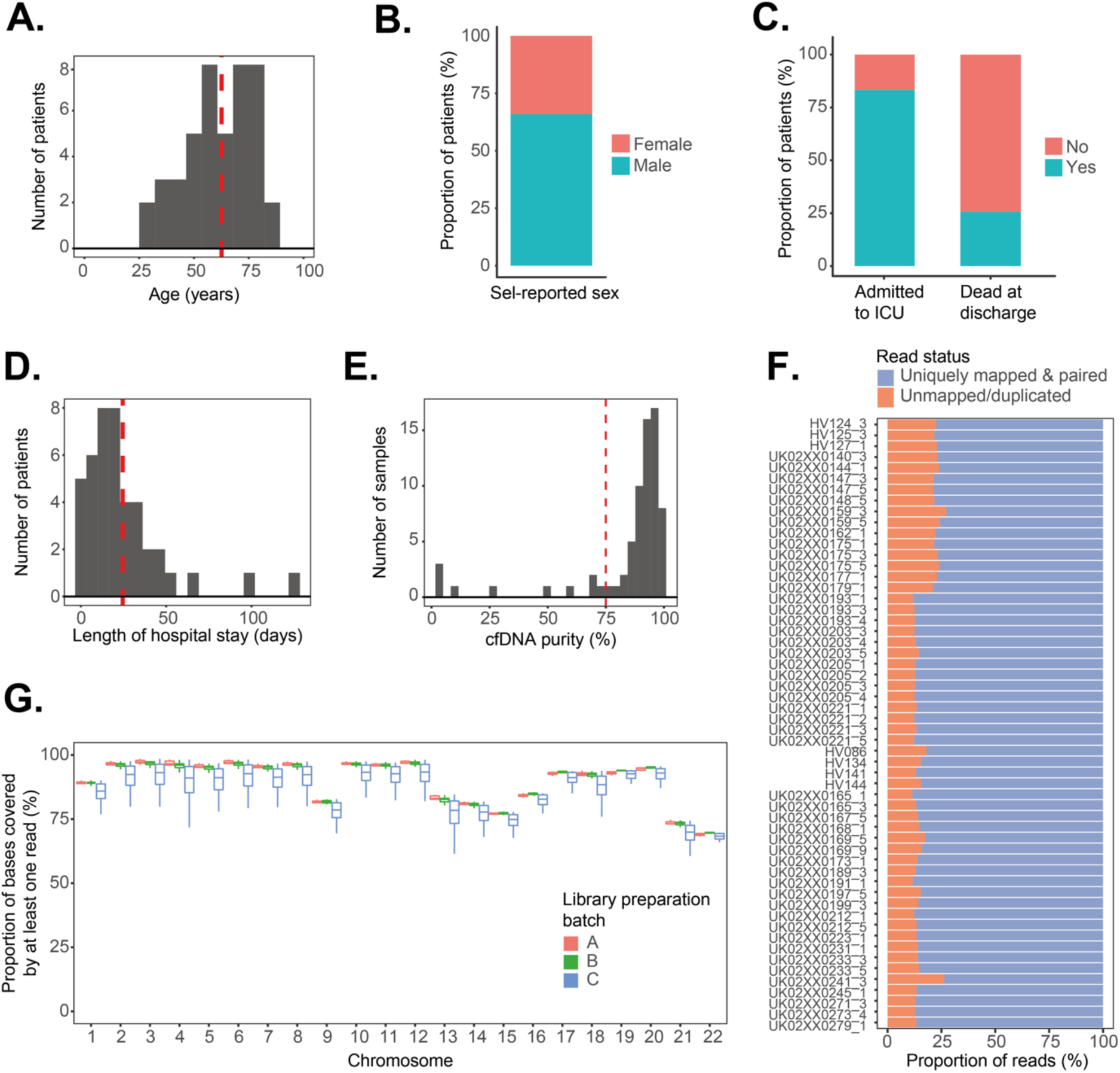
Cohort demographics and sample quality metrics. **A.** Distribution of self-reported age in our patient cohort (n=74 samples from 46 sepsis patients). The red dotted line indicates the mean age across patients. **B.** Distribution of self-reported sex in our patient cohort. Bar plots indicate patient proportions. **C.** Proportions of patients admitted to the ICU (left) and reported dead at hospital discharge (right). **D.** Distribution of hospital stay duration in our patient cohort. The red dotted line indicates the mean value. **E.** Distribution of cfDNA purity after isolation from plasma as estimated using capillary electrophoresis. The red dotted line indicates the cut-off 75% used for sample exclusion. **F.** Proportion of reads uniquely mapped to the human genome after sequencing in each sample in our study. **G.** Distribution of sequencing coverage in each chromosome is shown as the proportion of bases covered at > 1X (Y axis). Samples are stratified by TAPS library preparation batch. Box plots indicate the median and IQR for each sample.

**Figure S2.**
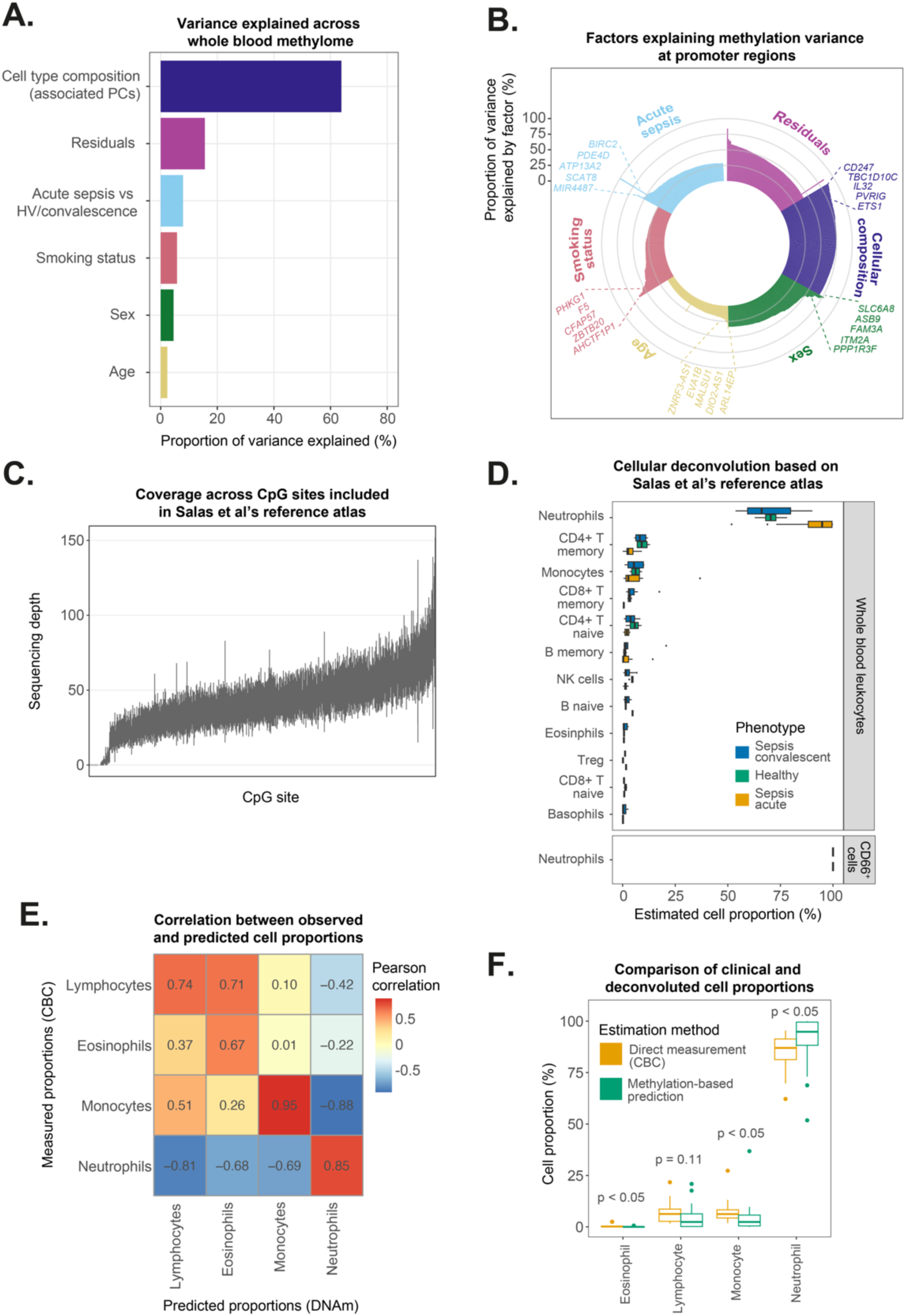
Analysis of the whole blood leukocyte methylome in 15 sepsis patients. **A.** The proportion of variance in WBL methylome (as profiled using EM-seq) (X axis) explained by different covariates (Y axis) was estimated using variance partitioning analysis. **B.** Variance partitioning estimates are shown for each CpG site included in the WBL methylation data set. The proportion of variance explained (Y axis) is shown for each assessed covariate (colour hue). Each bar represents a CpG site, with sites being ordered by decreasing variance estimate. **C.** Sequencing coverage (Y axis) achieved for each of the CpG sites included in Salas’s methylation tissue atlas (X axis). **D.** Cell type proportion estimates derived from WBL methylome deconvolution using EpiDISH and Salas’s methylation atlas. Colours indicate sample groups (n=15 acute sepsis patients, n=5 healthy controls, and n=9 convalescent/recovered sepsis patients). Bar plots indicate medians and IQRs for each cell type in the reference atlas. **E.** Correlation between cell proportions estimated from WBL methylome deconvolution and directly measured in a complete blood count (CBC) in hospital. Colours indicate Pearson correlation coefficients, with rows and columns ordered by similarity using hierarchical clustering. **F.** Cell proportion estimates (Y axis) derived from WBL methylome deconvolution and CBC measurements (colour hue). Box plots indicate median and IQR values for each cell type.

**Figure S3.**
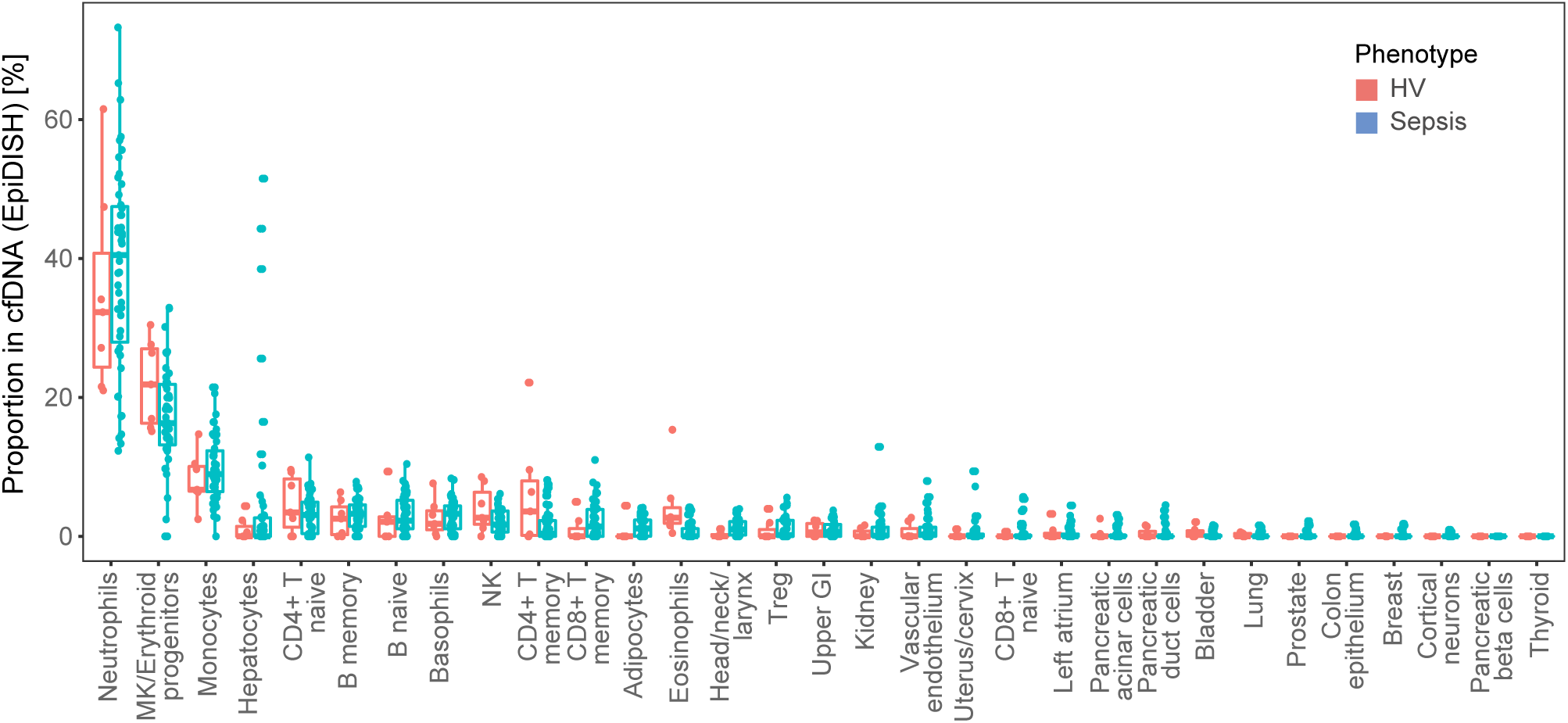
Cell-free DNA deconvolution based on Moss et al.’s methylation atlas. Proportion of cfDNA estimated to arise from different tissues based on methylome deconvolution using EpiDISH and the reference tissue atlas published by Moss et al. Box plots show median and interquartile ranges (IQR) of estimated proportions in sepsis patients (red) and healthy controls (blue).

**Figure S4.**
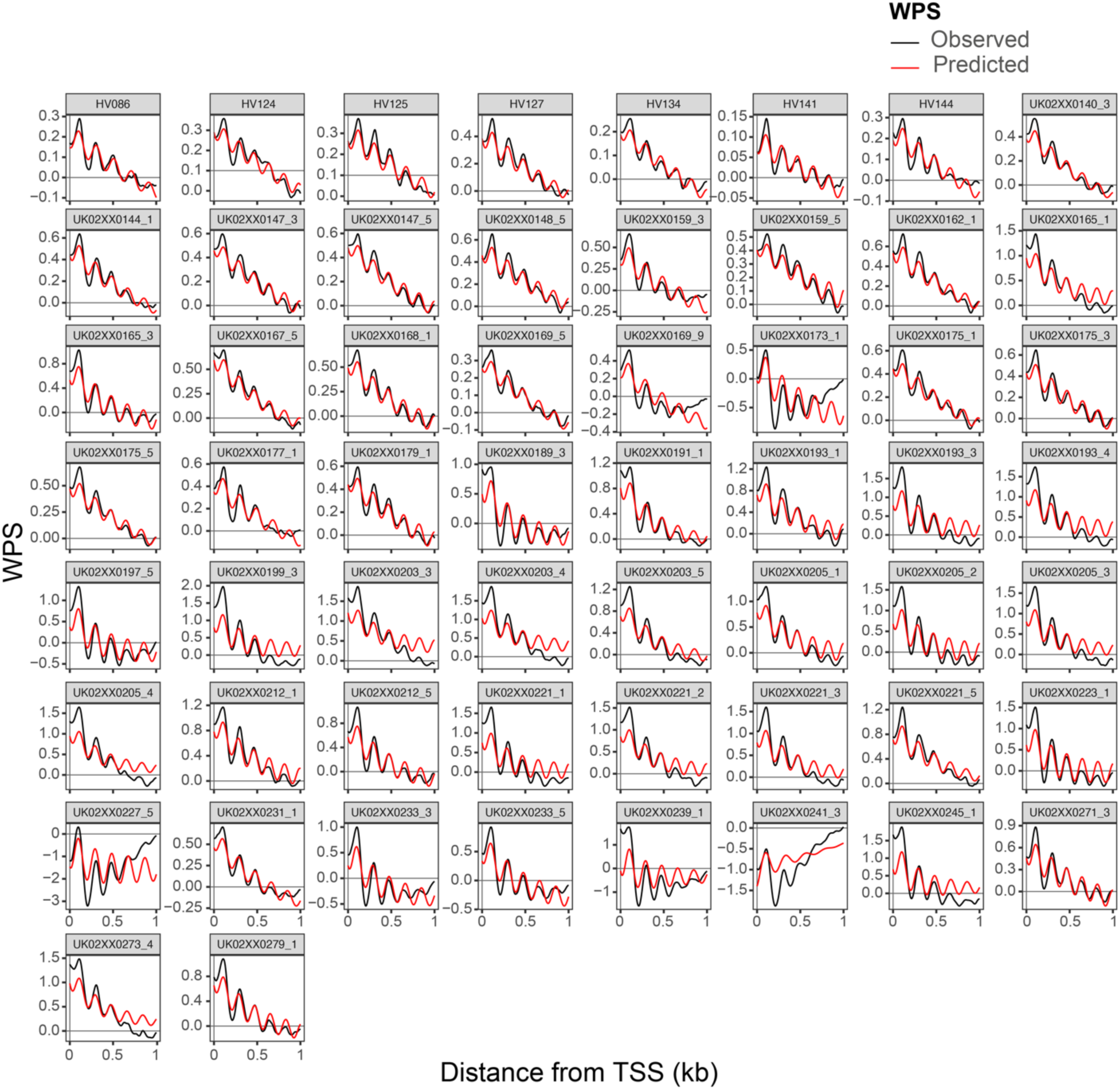
Dampened harmonic oscillator modelling of nucleosome phasing across samples. Observed WPS values at the TSS regions of all known genes (black lines) are shown alongside predictions derived from a dampened harmonic oscillator model (red lines) in each sample in our study. Model fitting was performed using non-linear least squares analysis. Each panel represents estimates from a different patient sample.

**Figure S5.**
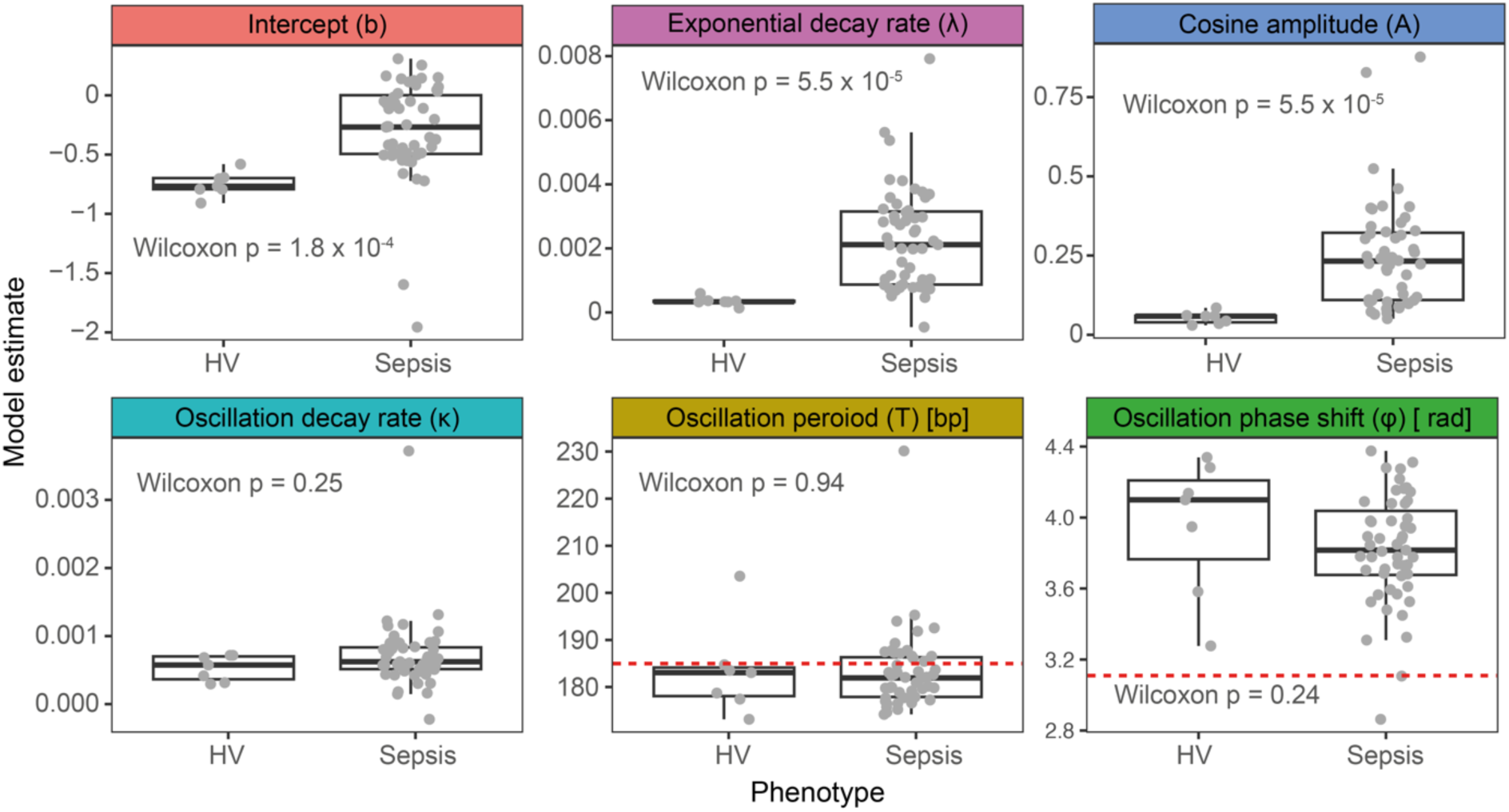
Nucleosome phasing parameter estimates. Parameter estimates from a dampened harmonic oscillator model were inferred using non-linear least squares analysis. Estimates (Y axis) are shown for each sample, stratified by disease status (X axis). Each panel shows estimates for a different parameter in the model equation. Dotted red lines indicate expected parameter values based on known biology. Box plots indicate median and IQRs for each parameter. P values were estimated using Wilcoxon rank sum tests.

**Figure S6.**
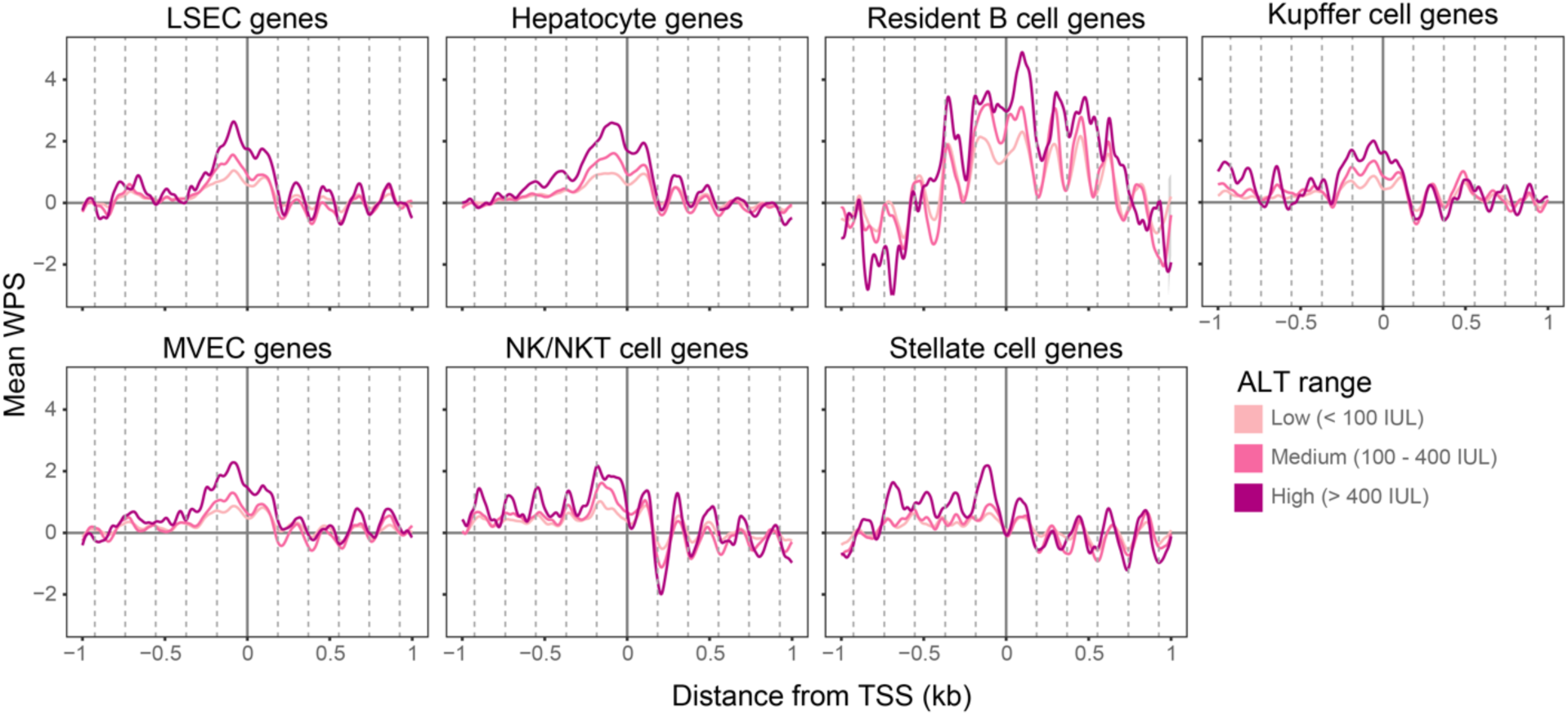
Nucleosome phasing strength at liver-specific gene sets correlates with liver dysfunction. Average WPS values at the TSS regions of genes specifically expressed in a variety of liver cell types, as determined by Aizarani et al.’s liver cell atlas. Lines indicate the average WPS values across samples classified as high, medium, or low levels of circulating ALT (colour hue). Dotted vertical lines indicate the expected position of nucleosomes around the TSS.

